# Molecular determinants of cotranslational chaperone recruitment to nascent p53

**DOI:** 10.64898/2026.04.17.719227

**Authors:** Laura Karpauskaite, Tomas B. Voisin, Sarah L. Maslen, Lois Kent, Grant A. Pellowe, J. Mark Skehel, David Balchin

## Abstract

Protein folding is fast relative to mRNA translation and nascent polypeptides begin to fold during their synthesis on the ribosome. The resulting cotranslational folding intermediates are accessible to diverse molecular chaperones that recognise incompletely folded client proteins. Here, we sought to understand how specific chaperone:client complexes are established during protein synthesis, using the chaperone-dependent tumor suppressor protein p53 as a model. By capturing interactomes at different points during p53 synthesis, we show that Hsp70, Hsp90 and TRiC bind competitively to nascent p53. While Hsp70 and Hsp90 bind promiscuously, TRiC discriminates between subtly different partially-folded states. TRiC recruitment is dictated by local cotranslational folding and exposure of specific sequence motifs, and this is tuned by cancer-associated mutations in the p53 DNA-binding domain. Nascent chain interactions with the ribosome surface disfavor TRiC binding, demonstrating that the ribosome imposes unique constraints on chaperone recruitment during protein synthesis. Our results establish molecular principles underlying chaperone prioritization during protein biogenesis.

## INTRODUCTION

Proteins begin to acquire their native structure as they are synthesised by the ribosome in a directional manner from the N-to C-terminus^1,2^. The stepwise process of protein synthesis allows the nascent chain (NC) to explore an evolving folding energy landscape that changes as each amino acid is added^3,4^. Although this can be beneficial (e.g. for folding multidomain proteins^5,6^), gradual cotranslational folding in the crowded cellular environment also creates a vulnerability. Partially-synthesised NCs lack complete sequence information required to fold, risking non-native intra-or intermolecular interactions^7^. Protein biogenesis is therefore critically safeguarded by molecular chaperones which bind NCs and guide their maturation^1,8^. The translating ribosome is a unique environment for chaperone action. The ribosome solubilizes and stabilizes folding intermediates^9–12,5,13,14^, and proximity to the ribosome surface may sterically restrict chaperone access. How these considerations shape chaperone recruitment to NCs is poorly understood.

Human cells express ∼150 cytonuclear chaperones, all of which may in principle participate in cotranslational folding of cytosolic proteins^1^. Early work showed that chaperones including TRiC, Hsp70 and Hsp40 act at the ribosome^8,15,16^, and subsequent studies have catalogued the NCs that interact with TRiC (yeast^17^ and human^18^), NAC (nematode^19^ and human^20^), Prefoldin (human^18^) and the fungi-specific Hsp70 homolog Ssb (yeast^17,21^). The binding of NAC and Ssb was shown to correlate with the emergence of short hydrophobic sequences from the ribosome^17,19–21^, whereas TRiC often binds later in synthesis when partial domains are exposed^17,18^. These studies demonstrate that cotranslational chaperone action is a pervasive feature of protein biogenesis, and imply that different chaperones form a cooperative network at the ribosome. The complete set of chaperones that sample different NCs remains to be determined, as does the molecular basis for their coordinated binding during cotranslational folding.

Chaperones are particularly important for the folding and function of conformationally-labile proteins, which constitute a substantial fraction of the human proteome^22^. An exemplar is the tumour suppressor p53, a transcription factor involved in regulating the cell cycle, DNA repair, apoptosis and senescence^23,24^. More than half of all cancers are associated with mutations in p53^25–27^. These most commonly occur in the conformationally unstable DNA-binding domain^28^, often destabilising it further, which can expose the hydrophobic core and lead to misfolding and aggregation^29^. Nascent p53 is therefore likely to be vulnerable to misfolding during synthesis, before collapse of the hydrophobic core. It remains unclear, however, how aggregation-prone segments of p53 are protected during cotranslational folding. p53 requires TRiC to fold efficiently^30^, and its activity is maintained via persistent conformational remodelling by Hsp70 and Hsp90^31,32^. Whether these chaperones also recognise folding intermediates of p53 on the ribosome is unknown.

Here, we develop a strategy to map the interactions of chaperones with nascent p53 in human cells, and rationalise their binding preferences. We show that chaperones recognise local features of cotranslational folding intermediates rather than global instability, and that interactions between NCs and the ribosome can mask chaperone binding sites to dictate chaperone recruitment. Our results establish principles underlying chaperone prioritization during protein biogenesis.

## RESULTS

### Cotranslational interactome of p53

During its synthesis on the ribosome, nascent p53 sequentially exposes different sequence and structural elements to the cytosolic chaperone network. We first sought to identify chaperones that interact with p53 at different stages of its biogenesis. To stabilise specific cotranslational folding intermediates, we used the XBP1u+ sequence to induce ribosome stalling in cells^5,33,34^. We chose stalling positions based on the domain architecture of p53, which consists of a DNA-binding (DBD) and a tetramerization (TET) domain flanked by flexible N-and C-termini (Fig. 1A). The resulting stalled ribosome:nascent chain complexes (RNCs) exposed either the unstructured N-terminal region of p53 (RNC_1-97_), part of the DBD (RNC_1-257_), the entire DBD (RNC_1-325_), or full-length p53 with a C-terminal 50 aa GlySer linker to extrude it from the ribosome (RNC_FL_) (Fig. 1B). As a negative control, we prepared a stalling construct containing only the GlySer linker (RNC_GS_). We expressed the constructs in Expi239F cells, isolated total ribosomes, then purified homogeneously stalled RNCs via an N-terminal 3xFLAG tag on the NC (Fig. 1C and S1A). We optimized cell lysis and RNC isolation steps to retain native interactors without resorting to crosslinking, allowing quantitative comparison between interactomes at different NC lengths. Importantly, this approach identifies NC-specific interactors rather than the total ribosome interactome as reported previously^35–38^.

**Fig. 1.**
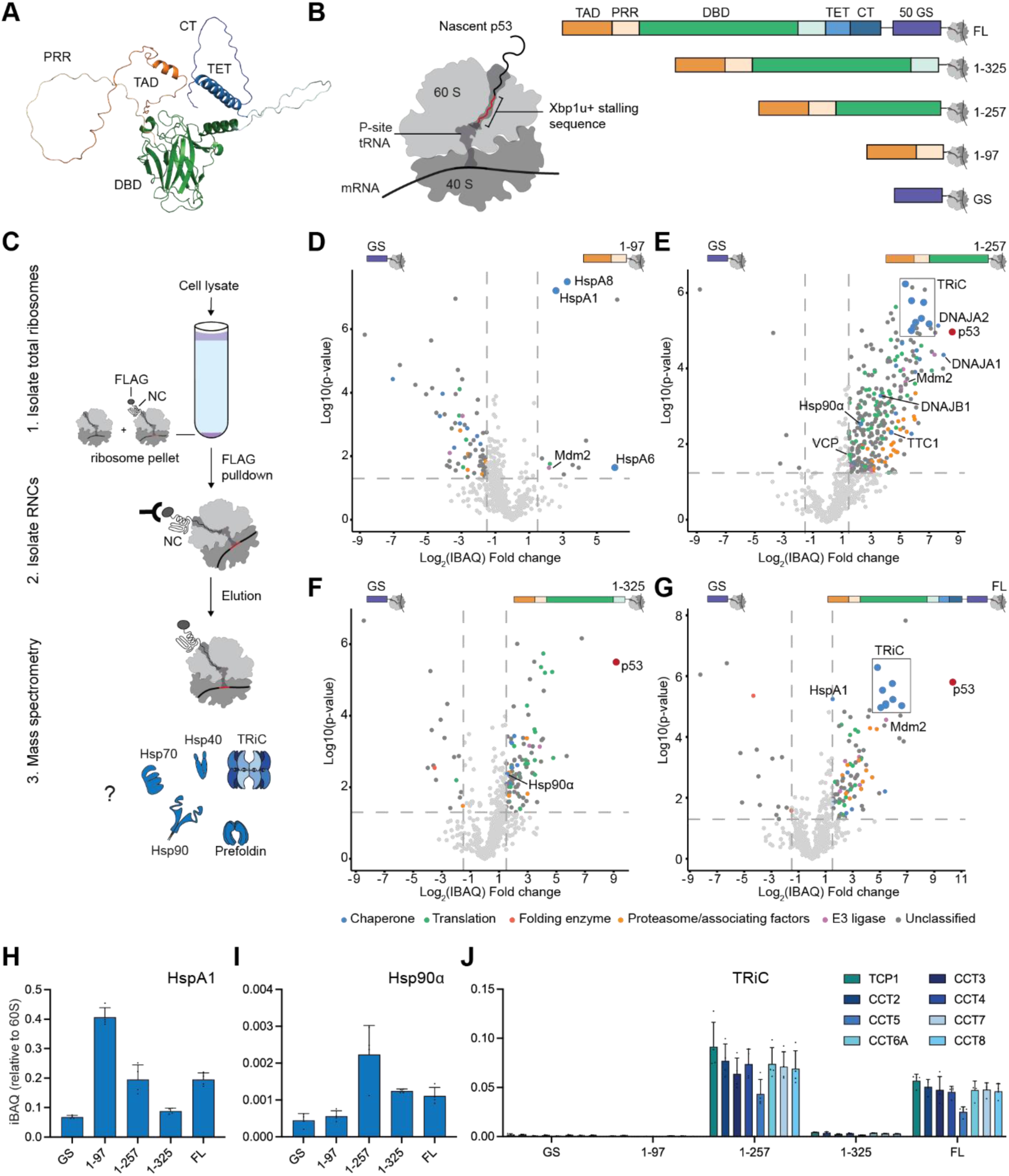
Cotranslational interactome of p53 (A) Predicted structure of p53 (AF-P04637-F1-v6), coloured by domain. TAD - transactivation domain (orange, residues 1-60); PRR - proline rich region (wheat, 61-93); DBD – DNA binding domain (green, 94-292); linker region (cyan, 293-325); TET – tetramerization domain (blue, 326-356); CT – C-terminal domain (teal, 357-393). (B) p53 ribosome:nascent chain complex (RNC) constructs. (C) Schematic of RNC isolation. (**D-G**) RNC interactomes. Volcano plots compare the interactome of RNC_GS_ with: (**D**) RNC_1-97_, (**E**) RNC_1-257_, (**F**) RNC_1-325_, or (**G**) RNC_FL_. Significant interactors (p-value>0.05, fold change>1.5) are annotated according to their function in proteostasis: chaperone (blue), translation (green), folding enzyme (red), E3 ligase (pink), proteasome and associated factors (orange), unclassified (grey). The bait NC is coloured red. (**H-J**) Quantification of chaperone occupancy. Intensity-based absolute quantification (iBAQ) of (**H**) HspA1, (**I**) Hsp90α, and (**J**) TRiC subunits, normalised to the average iBAQ of 60S ribosomal proteins in each sample. Error bars represent SD, n=4 biological replicates. See also Fig S1 and Data S1.

Mass spectrometry confirmed that the RNCs contained all 80S ribosomal proteins at stoichiometric levels, and revealed additional interactors specific to each NC length (Fig. 1D-G, S1B,C and Data S1). Chaperones were enriched in all p53 RNCs compared to RNC_GS_, but their identity and relative abundance varied. Several Hsp70s (HspA1, HspA6, HspA8) were abundant in all RNCs (>10% of RNC occupancy) but most strongly enriched in RNC_1-97_ (∼40% occupancy for HspA1), indicating that they are preferentially recruited early in synthesis when only the disordered N-terminus of p53 is exposed (Fig. 1H and S1D,E). Specific J-domain proteins (JDPs: DNAJA1, DNAJA2, DNAJB1) were enriched only in partial DBD RNC_1-257_ (Fig. 1E). DnaJA1/2 were present at ∼10-fold higher levels than DNAJB1, consistent with the recent finding that A-class JDPs preferentially bind the destabilised p53 DBD independently of Hsp70^39^ (Fig. S1F,G). Both Hsp90β (Hsp90AB1) and Hsp90α (Hsp90AA1) copurified with RNCs in low amounts (∼0.2-0.5% occupancy) (Fig. 1I and S1H). Hsp90β was slightly more abundant overall, but only Hsp90α was significantly enriched in RNC_1-257_ and RNC_1-325_ compared to RNC_GS_. We also detected several Hsp90 cochaperones, but only TTC1/TPR1 was consistently enriched, in RNC_1-257_ (Fig. 1E). The TRiC chaperonin complex (all 8 subunits at equivalent levels, ∼5-10% occupancy) was strongly enriched in both RNC_1-257_ and RNC_FL_ (Fig. 1J). We detected small amounts of the TRiC cochaperone prefoldin, but it was not significantly enriched, suggesting that prefoldin does not stably bind nascent p53 concomitantly with TRiC (Fig. S1J). The ribosome-binding factor NAC (NACA and BTF3) was present but not enriched over the control (Fig. S1I). We did not detect subunits of mRAC (HspA14 and DnaJC2).

In addition to chaperones, several protein quality control factors were enriched in p53 RNCs (Fig. 1D,E,G). Proteasome subunits and VCP were enriched in RNC_1-257_ which exposes the partial DBD, and the p53-specific E3 ligase Mdm2^40,41^ was enriched in all RNCs except RNC_1-325_ which exposes the complete DBD. The selective enrichment of quality control factors at specific NC lengths suggested that they are recruited based on features of the NC rather than ribosome stalling. Indeed, ribosome-associated quality control factors (e.g. NEMF and Ltn1^42^) were not detected.

These data establish the pattern of chaperone engagement during p53 synthesis on the ribosome. Early in synthesis, the disordered N-terminus recruits Hsp70s but no other chaperones. Subsequent synthesis of the partial DBD results in a folding intermediate which recruits TRiC, Hsp90 and JDPs, at the expense of Hsp70s. Completion of DBD synthesis then disfavours binding of all chaperones except Hsp90α. Upon emergence of full-length p53 from the ribosome, TRiC is selectively recruited once more, albeit in a smaller amount than to the partial DBD. Quantitative comparison of chaperone occupancies suggests that TRiC and Hsp70 bind relatively stably to specific p53 RNCs, while Hsp90 binding is more transient.

### Chaperone coordination during p53 synthesis

Hsp70, Hsp90 and TRiC were all enriched on RNC_1-257_ exposing the incomplete p53 DBD, and we next sought to understand how their binding is coordinated. The complete DBD folds into an immunoglobulin-like β-sandwich with strands β1-10 flanked by helices α1 and α2 (Fig. 2A). We purified a series of RNCs sampling sequential intermediates during DBD synthesis, and probed for chaperones by immunoblotting (Fig. 2B). While levels of Hsp70 and Hsp90β did not vary substantially during DBD synthesis, TRiC was highly selective. TRiC was recruited just prior to the emergence of β5 from the exit tunnel (RNC_1-197_), i.e. ∼halfway through DBD synthesis, and disengaged when all β-strands were available for folding (RNC_1-296_).

**Fig. 2.**
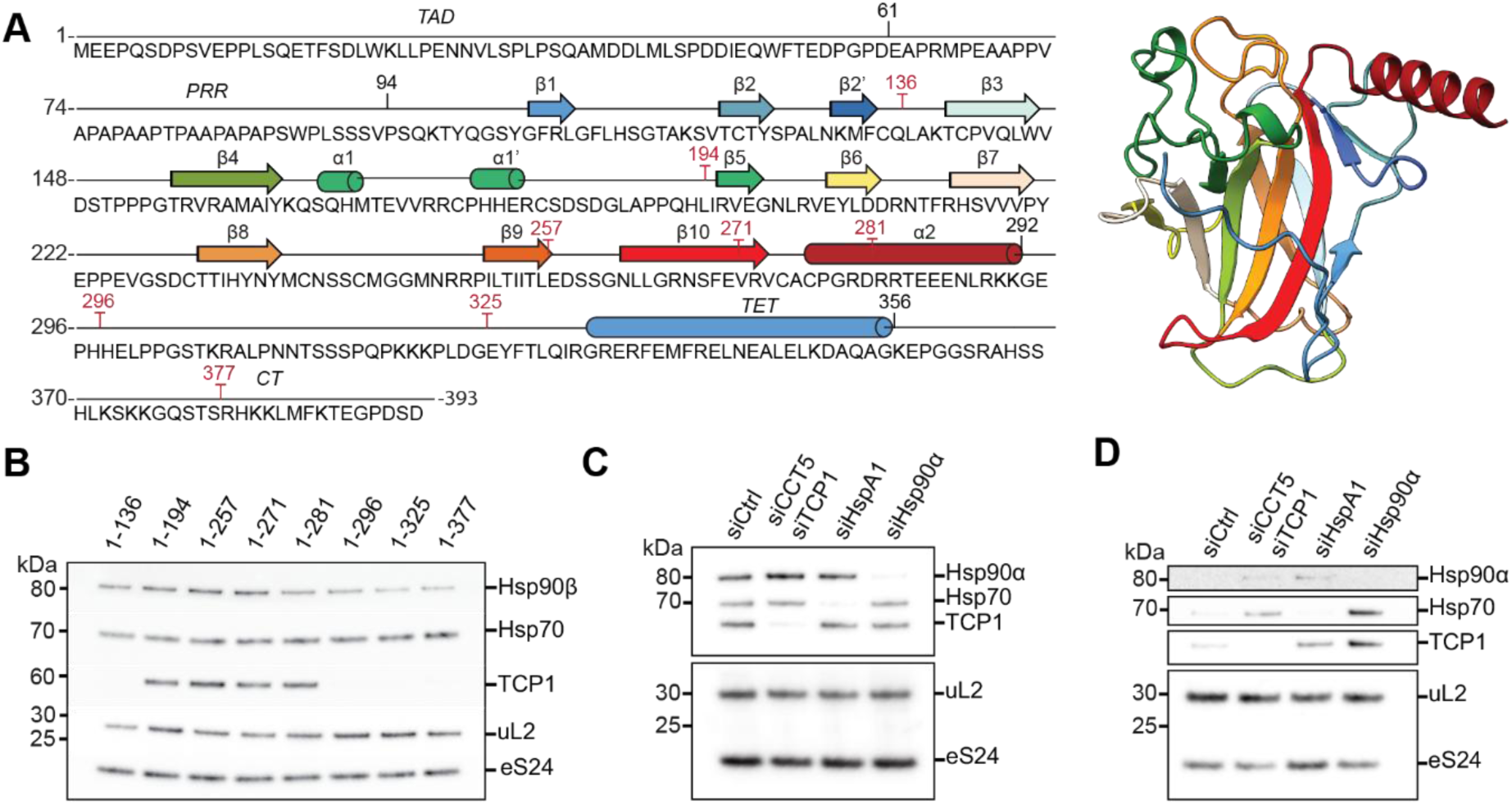
Chaperone coordination during p53 synthesis (A) Left: sequence and secondary structure of p53 with the DBD coloured in a rainbow gradient from N to C terminus. Stalling positions are indicated in red. Right: predicted structure of the DBD (residues 94-293, AF-P04637-F1-v6). (B) Chaperone recruitment to the nascent DBD. RNCs stalled during DBD synthesis were purified and analysed by immunoblotting for the indicated proteins. (**C-D**) Effect of chaperone knockdown. Cells were treated with control siRNA (siCtrl), or siRNA targeting TRiC subunits (TCP1/CCT5), Hsp70 (HspA1), or Hsp90α, before expression of RNC_1-257_. Immunoblots of the total ribosome fraction (**C**) and purified RNC (**D**) are shown. See also Fig S2.

To test whether TRiC, Hsp70 and Hsp90 cooperate or compete to bind the DBD, we used siRNA to knock down each chaperone individually. A combination of siRNAs targeting TRiC subunits (CCT1 and CCT5) was highly efficient in depleting TRiC in cells^43^, but total levels of Hsp90α and Hsp70 were only partially reduced by the corresponding siRNAs (Fig. S2A,D). However, Hsp90α and Hsp70 were near-completely depleted in the ribosomal fraction when silenced by their respective siRNAs (Fig. 2C, S2B). Probing for chaperones on purified RNC_1-257_ revealed competitive binding: depletion of any chaperone increased levels of the other two chaperones on the RNC (Fig. 2D and S2C,E,F). This effect was particularly pronounced for the Hsp90α knockdown, which led to strong enrichment of Hsp70 and TRiC. Thus, although Hsp90α does not bind stably to the NC (c.f. chaperone occupancy measured by MS, Fig. 1I), it regulates the access of other chaperones.

In summary, we find that Hsp70 and Hsp90 dynamically sample the p53 DBD throughout its synthesis, while TRiC acts in a narrower window but binds more stably. TRiC selects a folding intermediate that forms upon exposure of approximately half of the DBD from the ribosome, and is outcompeted by Hsp90 only when the DBD is complete.

### TRiC discriminates between folding-compromised NCs

We next sought to characterise the determinants of TRiC binding to nascent p53. First, we tested the effect of nucleotides on TRiC binding to RNC_1-257_. Consistent with a bona fide TRiC:client interaction^44–46^, copurifying TRiC was depleted by ATP (which promotes client release), stabilised by ADP, and partially retained by trapping the transition state of ATP hydrolysis using ATP/AlF_X_ (Fig. 3A). We next examined structural features of the NC. TRiC disengages nascent p53 upon completion of DBD synthesis, which we hypothesised was due to cotranslational folding of the DBD. To test this, we probed the conformation of ribosome-tethered p53 by limited proteolysis (Fig. S3A). While RNC_1-257_ exposing the partial DBD was rapidly degraded by Proteinase K, proteolysis of RNC_1-325_ exposing the complete DBD resulted in the accumulation of a ∼60 kDa intermediate. The intermediate was not detected by an antibody raised against the N-terminus of p53 (DO1, epitope residues 20-26), indicating that it had lost the flexible N-terminus. Furthermore, it shifted in size to ∼35 kDa upon RNase A treatment, showing that it was still attached to the ribosome via peptidyl-tRNA. The ∼35 kDa intermediate is diagnostic for native folding, since it also accumulated during proteolysis of FL p53. As an orthogonal probe of folding, we used conformation-specific p53 antibodies to immunoprecipitate RNCs. Pab1620 recognises surface features of folded p53, whereas Pab240 recognises a buried epitope that is exposed upon unfolding^47–49^. As Pab1620 does not bind monomeric p53^50^ (Fig. S3B,C), we induced assembly by fusing the normally C-terminal TET to the N-terminus of RNC_1-257_ and RNC_1-325_. RNC_TET-1-257_ was immunoprecipitated only by Pab240, while RNC_TET-1-325_ was immunoprecipitated by Pab1620, indicating that the DBD adopts a native-like conformation in RNC_TET-1-325_ but not RNC_TET-1-257_ (Fig. S3D-F)_._ Together, these data show that completion of DBD synthesis triggers cotranslational folding close to the ribosome surface, coinciding with loss of affinity for TRiC.

**Fig. 3.**
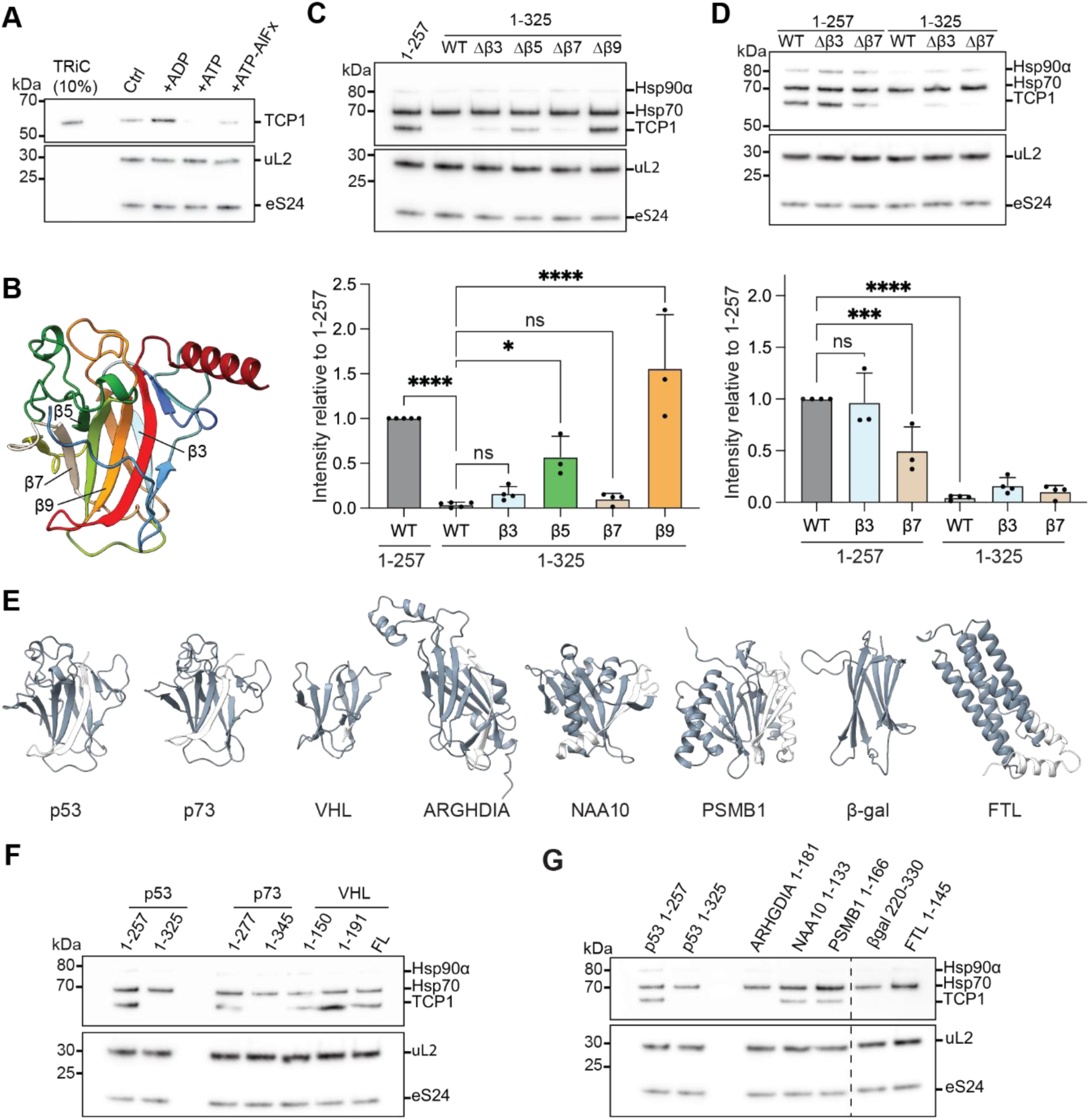
TRiC discriminates between folding-compromised NCs (A) Effect of nucleotides on TRiC recruitment. RNC_1-257_ was purified either without additional nucleotides (Ctrl), or with additional 1 mM ADP, 1 mM ATP, or 1 mM ATP-AlFx (1 mM ATP, 6 mM NaF, 1 mM Al(NO_3_)_3_). TRiC (TCP1) was probed by immunoblotting. Purified TRiC at 10% of the RNC amount is shown for reference. (B) Predicted structure of p53 DBD (residues 94-293, AF-P04637-F1-v6) highlighting β3, β5, β7, and β9. (C) Chaperone recruitment to DBD with β-strand deletions. Top: WT RNC_1-257_, and RNC_1-325_ with different β-strands deleted, were purified and analyzed by immunoblotting for the indicated proteins. Bottom: quantification of TCP-1 signal, normalised to the loading control (uL2) and then to WT RNC_1-257_. Data represent mean ± SD, n=3-5 biological replicates. Statistical analysis was performed using one-way ANOVA with Dunnett’s post hoc test. Significance relative to control is indicated as p >0.05 (ns), p < 0.05 (*), p < 0.01 (**), p < 0.001 (***), p < 0.0001 (****). (D) As in (**C**), for variants of RNC_1-257_ with β-strands deleted. Data represent mean ± SD, n=3-4 biological replicates (E) Structures of p53 DBD (AlphaFold3), p73 (PDB:3VD0), VHL (PDB:1LM8), ARHGDIA (AlphaFold3), NAA10 (PDB:6PPL), PSMB1 (PDB:5GJQ), β-galactosidase domain 2 (β-gal, AlphaFold3), and FTL (AlphaFold3). The region included in each RNC construct is coloured slate. (F) Chaperone recruitment to β-sandwich RNCs. RNCs with the indicated residue numbering were purified and analyzed by immunoblotting. (G) Chaperone recruitment to β-strand-rich RNCs. RNCs with the indicated residue numbering were purified and analyzed by immunoblotting. See also Fig S3 and S4.

Since TRiC binds the partially-synthesised DBD lacking one or more C-terminal β-strands (Fig. 2B), we wondered whether the absence of other β-strands from the DBD β-sandwich would suffice to recruit TRiC. To test this, we deleted β3, β5, β7 or β9 from RNC_1-325_ exposing the complete DBD (Fig. 3B,C). Although all variants were structurally destabilised as shown by limited proteolysis (Fig. S4A), only Δβ5 and Δβ9 recruited TRiC. We hypothesised that β3/β7 deletions may not recruit TRiC because these β-strands harbour TRiC binding sites. To test this, we deleted β3 or β7 from RNC_1-257_ which stably binds TRiC (Fig. 3D). While β3 deletion had no effect, β7 deletion reduced copurifying TRiC, indicating that β7 contains a TRiC binding site. TRiC therefore recognises DBD variants lacking β5 or β9, fails to recognise a variant lacking β3, and interacts with β7. In contrast to TRiC, neither Hsp70 nor Hsp90 strongly discriminated between Δβ variants, consistent with their recruitment throughout DBD synthesis (Fig. 2B and S4B,C).

These data indicate that TRiC does not bind partially-folded NCs indiscriminately. To understand if TRiC recognises sequence or structural features of the incomplete DBD, we tested 6 other proteins with β-strand-rich domains (Fig. 3E). At two extremes of sequence identity were human p73 (63% sequence identity to p53 in the DBD^51^) and domain 2 of *E. coli* β-gal (9% sequence identity but high structural similarity with 7 Å RMSD to the DBD). We designed each RNC to expose an incomplete domain on the ribosome that mimics p53 RNC_1-257_. An all-α-helical protein FTL (1-145) was included for comparison. We found that p73, VHL, NAA10 and PSMB1 recruited TRiC, whereas ARHGDIA, β-gal, and FTL did not (Fig. 3F,G). Thus, neither domain structure nor β-strand content fully explain TRiC binding preferences. Interestingly, unlike p53 and p73, VHL strongly recruited TRiC even when the complete β-sandwich was exposed on the ribosome (Fig. 3F). This is likely due to instability of VHL in the absence of Elongin B/C, and is consistent with a role for TRiC in assembling the VHL/Elongin complex^52^.

In summary, although conformational instability is a prerequisite for TRiC binding, neither this property nor β-strand propensity is definitive. Instead, our data argue that TRiC binding is driven by exposure of specific sequence motifs in the nascent polypeptide.

### Cancer-associated mutations tune chaperone recruitment

We next considered whether more subtle sequence variations influence cotranslational chaperone recruitment. p53 is mutated in >50% of cancers, with most mutations compromising tumour suppressor function by destabilising the DBD^28^. To understand whether chaperones distinguish between WT and mutant conformations of cotranslational folding intermediates, we introduced cancer-associated mutations into p53 RNCs. Importantly, NCs are solubilized by tethering to the ribosome^53^, allowing us to test even highly destabilizing sequence variants that are normally experimentally inaccessible due to protein aggregation.

We first sought to induce extreme destabilisation by combining multiple mutations^54^ to produce two variants: V1 (R175H, Y220S, Y236C, R248Q) and V2 (Y163S, Y220S, Y236C, R249S)

(Fig. 4A). We found that the effect of the mutations on chaperone binding was dependent on NC length. In RNC_1-325_ with a complete DBD, neither V1 nor V2 increased TRiC recruitment (Fig. 4B). In contrast, both sets of mutations decreased TRiC binding in the context of the incomplete DBD (RNC_1-257_) (Fig. 4B). Unlike TRiC, Hsp70 and Hsp90 levels were not significantly affected (Fig. S5A,B). Thus, severe thermodynamic destabilization of a folded domain is insufficient to trigger TRiC recruitment to NCs, and TRiC binding to a partially-synthesised NC can be impaired by mutations that destabilize the native state.

**Fig. 4.**
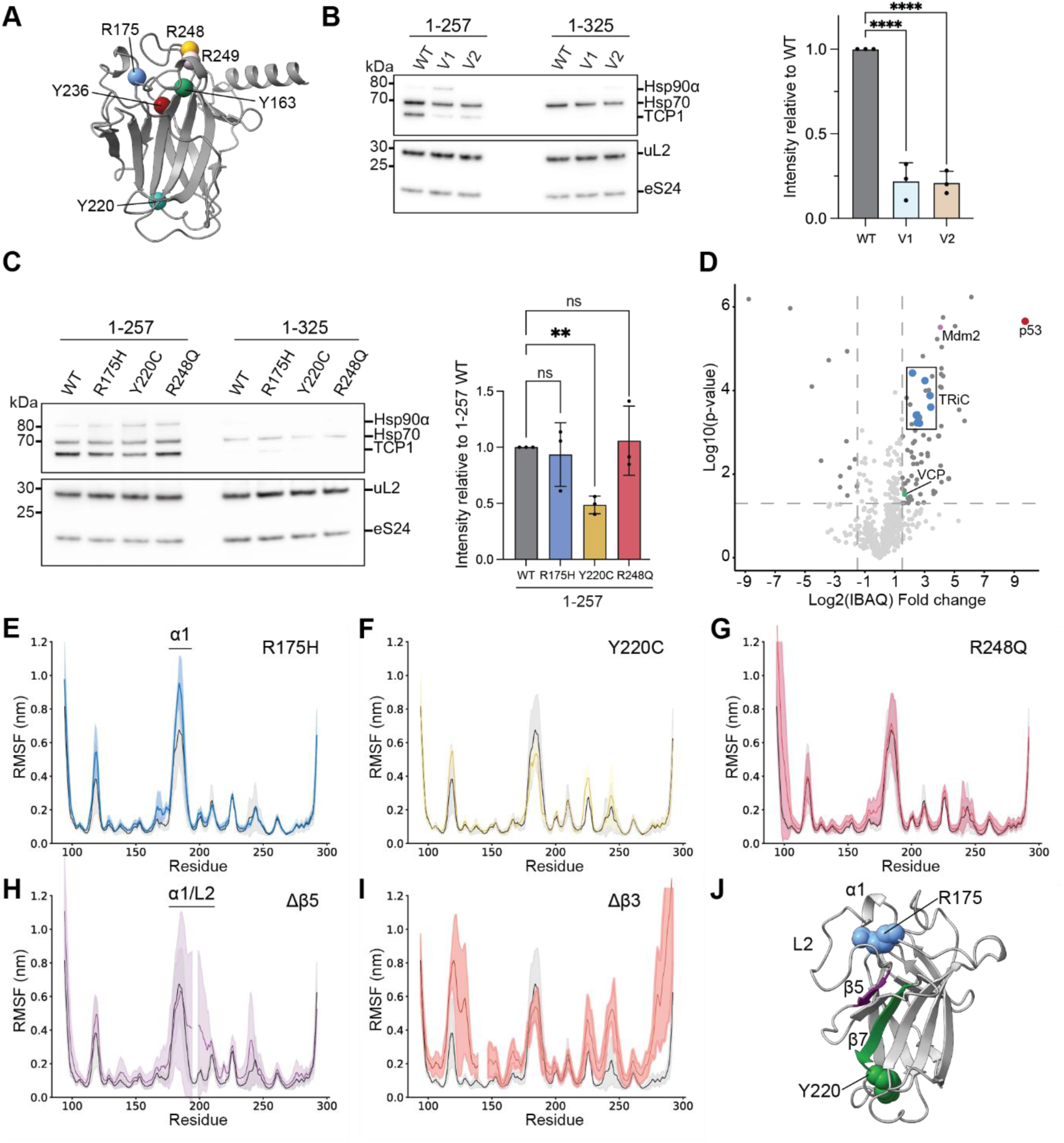
Cancer-associated mutations tune chaperone recruitment (A) Predicted structure of p53 DBD (residues 94-293, AF-P04637-F1-v6) with mutation sites highlighted. (B) Chaperone recruitment to RNCs with combinations of mutations. Left: RNC_1-257_ or RNC_1-325_ with V1 (R175H, Y220S, Y236C, R248Q) or V2 (Y163S, Y220S, Y236C, R249S) mutations were purified and analyzed by immunoblotting. Right: quantification of TCP-1 signal for RNC_1-257_, normalised to the loading control (uL2) and then to WT. Data represent mean ± SD, n=3 biological replicates. (C) Chaperone recruitment to RNCs with point mutations. Left: RNC_1-257_ or RNC_1-325_ with the indicated mutations were purified and analyzed by immunoblotting. Right: quantification of TCP-1 signal for RNC_1-257_, normalised to the loading control (uL2) and then to WT. Data represent mean ± SD, n=3 biological replicates. Statistical analysis was performed using one-way ANOVA with Dunnett’s post hoc test. Significance relative to control is indicated as p >0.05 (ns), p < 0.05 (*), p < 0.01 (**), p < 0.001 (***), p < 0.0001 (****). (D) Volcano plot comparing the interactomes of RNC_GS_ and RNC_1-325_ R175H. Significantly enriched proteins (p-value>0.05, fold change>1.5) are indicated. (**E-I**) Dynamics of mutant DBDs. Per-residue root-mean-square fluctuation (RMSF) of variant (coloured lines) and WT (black) p53 DBDs from molecular dynamics simulations. Solid lines: average RMSF; shaded intervals: SD, n=3 independent simulations. (**J**) Structure of p53 DBD, highlighting regions (α1, L2) and perturbations (R175 mutation, β5 deletion) that control the accessibility of chaperone binding motifs such as β7/Y220. See also Fig S5.

We next tested the effects of single point mutations on chaperone recruitment. We focused on three frequently-occurring cancer-associated mutations^28^ that are spatially distributed in the DBD: R175H, Y220C and R248Q (Fig. 4A). While Hsp70 and Hsp90 were unaffected by mutation of RNC_1-325_ or RNC_1-257_, TRiC binding was sensitive to point mutations (Fig. 4C and S5C,D). In particular, Y220C substantially reduced TRiC binding to RNC_1-257_. Y220 is located in a loop directly following β7, which we found to mediate TRiC recruitment (Fig. 3D). These data suggest that a sequence element encompassing β7 and Y220 is recognized by TRiC when exposed in the partially-synthesized DBD of p53.

Although the effect was small, we noticed an increase in TRiC recruitment to RNC_1-325_ R175H (Fig. 4C). To validate this finding, and to identify other factors that might recognise the destabilized NC, we used MS to determine the interactome of RNC_1-325_ R175H (Fig. 4D). These data confirmed the enrichment of TRiC, but not Hsp70 or Hsp90, on the mutant RNC. DnaJA1 levels also increased, consistent with the recent observation that it recognises the destabilized DBD^39^ (Fig. S5E). Moreover, RNC_1-325_ R175H recruited quality control factors including Mdm2 and VCP, which we also detected in wild-type RNC_1-257_.

We were intrigued by the finding that, among different destabilizing mutations, only R175H increased TRiC binding to the p53 DBD. To probe structural features that correlate with chaperone binding, we performed all-atom molecular dynamics simulations of WT and mutant p53 DBDs off the ribosome. None of the mutations increased the radius of gyration or solvent-accessible surface area of the DBD, suggesting that global properties of the DBD do not determine chaperone recruitment (Fig. S5F,G). However, per-residue root mean square fluctuation (RMSF) revealed increased dynamics in α1 that were specific to R175H, suggesting that local instability in this region may be important for TRiC recognition (Fig 4E-G). We wondered whether the same phenomenon would explain the difference in TRiC recruitment between β-strand deletions in the DBD (Fig. 3C). Indeed, Δβ5 which recruits TRiC showed a pronounced increase in RMSF of α1 and the adjacent L2 (Fig. 4H). In contrast, Δβ3 which does not recruit TRiC showed increased dynamics in β1,2,8,10 and α2, but not α1/L2 (Fig. 4I).

Together, our data suggest that TRiC binding is driven by specific sequence motifs - including, but not limited to, β7 and Y220 - that are exposed during DBD synthesis or unmasked by destabilisation of α1/L2 (Fig. 4J). As a result, the effect of cancer-associated mutations on chaperone recruitment depends on their location rather than the degree to which they are destabilizing. Only TRiC (not Hsp70 or Hsp90) was sensitive to NC mutations, indicating that even single point-mutations can tip the balance between different chaperone-directed maturation pathways.

### Cotranslational assembly of p53 recruits TRiC to the ribosome

We next turned our attention to the recruitment of TRiC to RNC_FL_. TRiC binding to RNC_FL_ was surprising, since TRiC disengages p53 upon DBD folding in RNC_1-325_ (Fig. 2B). In addition to a complete DBD as in RNC_1-325_, RNC_FL_ also contains the TET oligomerisation domain and a C-terminal GlySer linker (Fig. 1B). The linker did not mediate TRiC binding, since appending the same linker to RNC_1-325_ did not recruit TRiC (Fig. 5A). Nor is TET itself a TRiC client, since TRiC did not bind an RNC exposing TET alone (Fig. 5B). We also considered whether the C-terminal TET and linker may promote TRiC binding by allosterically destabilising the DBD. We used hydrogen/deuterium exchange-mass spectrometry (HDX-MS) to compare the conformational dynamics of isolated off-ribosome FL-p53 and a truncation consisting of residues 1-325 (Fig. S6A). Deuterium uptake in the DBD was near-identical between the two proteins (ΔDa < 0.5), indicating that the DBD is not destabilised by synthesis of C-terminal regions.

**Fig. 5.**
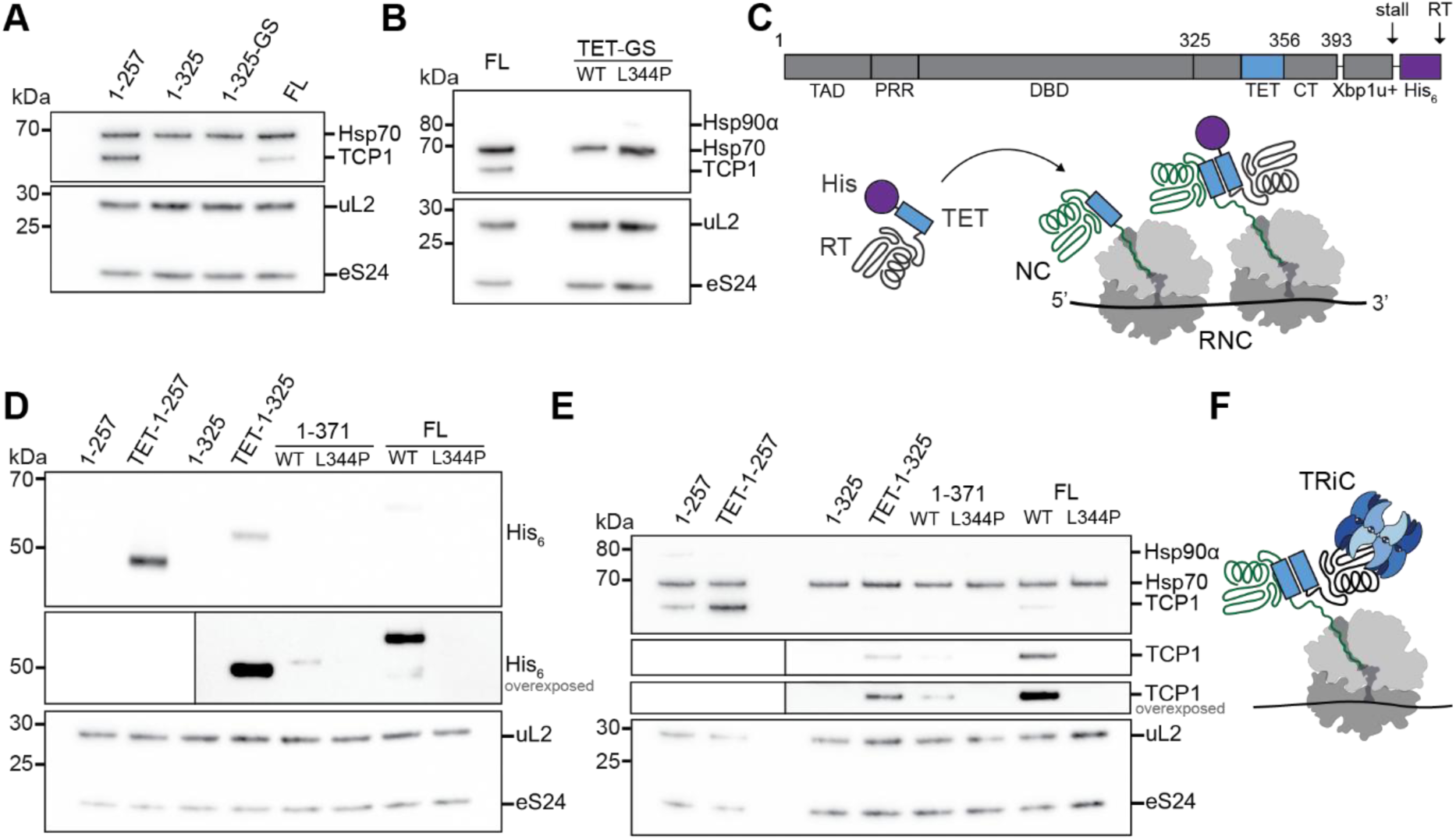
Cotranslational assembly of p53 recruits TRiC to the ribosome (A) C-terminal linker does not recruit TRiC to the nascent DBD. Purified RNCs were analyzed by immunoblotting. 1-325-GS contains a 50 aa Gly-Ser linker between the p53 fragment and stalling sequence. (B) TET does not bind TRiC. Purified RNCs were analyzed by immunoblotting. RNC_TET-GS_ consists of the TET domain and a C-terminal 50 aa Gly-Ser linker to extrude it from the ribosome. The L344P mutation prevents TET oligomerization. (C) Schematic of experiment to test cotranslational assembly. Ribosomes stochastically translate through XBP1u+, resulting in accumulation of a read-through product (RT) with a His_6_ tag in addition to the stalled RNC. TET-dependent recruitment of His_6_-tagged species to RNCs indicates cotranslational assembly. (D) Released p53 assembles with NCs via the TET domain. Purified RNCs were analyzed by immunoblotting for His_6_ to detect co-assembling RT product. In TET-1-257 and TET-1-325, TET is fused to the N-terminus of the NC. The L344P mutation prevents TET oligomerization. Where indicated, the immunoblot was cut and overexposed. (E) TRiC recruitment correlates with cotranslational assembly. Purified RNCs as in (**D**) were analyzed by immunoblotting for chaperones. Where indicated, the TCP1 (TRiC) immunoblot was cut and re-imaged to strengthen the signal, and overexposed. (F) Model of TRiC recruitment to RNCs exposing TET. TRiC binds mature p53 off the ribosome, and is indirectly recruited to nascent p53 by cotranslational assembly. See also Fig S6.

Since NC sequence features did not explain TRiC recruitment to RNC_FL_, we considered a role for cotranslational p53 assembly. p53 forms stable dimers^55^, and tetramerises in complex with cognate DNA^56–58^. p53 has previously been shown to assemble into homodimers during translation^59^, and assembly is in principle possible in RNC_1-371_ and RNC_FL_ which expose the TET domain (Fig. 2A). To detect assembly, we exploited the fact that ribosomes occasionally “read through” the XBP1u+ stalling sequence to produce a full-length protein alongside the stalled RNCs. The read-through product was distinguished from the NC by a His_6_ tag placed between XBP1u+ and the stop codon (Fig. 5C). Consistent with cotranslational assembly via TET, RNC_1-371_ and RNC_FL_ co-purified with His-positive species, whereas RNC_1-257_ and RNC_1-325_ did not (Fig. 5D). Co-purification of p53-His_6_ was abolished by the L344P mutation which disrupts TET^60^, and could be induced in RNC_1-257_ and RNC_1-325_ by fusing TET to the N-terminus (Fig. 5D). Ribosome-tethered p53 exposing TET can therefore assemble with a released partner. Furthermore, assembly correlated with TRiC recruitment. The L344P mutation blocked TRiC binding to RNC_1-371_ and RNC_FL_, while TRiC was recruited in increased amounts when TET was appended to RNC_1-257_ and RNC_1-325_ (Fig. 5E). Unlike TRiC, Hsp70 and Hsp90α were insensitive to the assembly status of RNCs.

The correlation between TRiC recruitment and p53 assembly competence could be explained if TRiC preferentially binds assembled p53. To test this, we expressed non-stalled FLAG-tagged FL-p53 WT, an assembly-incompetent point mutant (L344P), and an assembly-incompetent truncation lacking TET (1-325) in Expi293F cells. FLAG pulldown recovered TRiC in all cases, demonstrating that TRiC binds both monomeric and oligomeric p53 post translation (Fig. S6B). Thus, although TRiC can recognise monomeric p53 with a complete DBD, it fails do so when p53 is tethered to the ribosome. We conclude that TRiC is recruited to RNC_FL_ by an interaction with the fully-translated co-assembling subunit, rather than with the ribosome-tethered NC itself (Fig. 5F).

To test whether TRiC influences the efficiency of cotranslational assembly, we quantified recruitment of p53-His_6_ to RNC_FL_ following TRiC knockdown (Fig. S6C-E). Co-assembling p53 was increased when TRiC was depleted, indicating that TRiC impedes cotranslational assembly, possibly by sterically interfering with TET oligomerization.

### Nascent p53 escapes TRiC by binding the ribosome

TRiC binds FL-p53 off the ribosome, but ignores ribosome-tethered p53 once the DBD is complete. These findings suggest that the ribosome influences the conformation or accessibility of nascent p53 to disfavour TRiC binding. To test this, we characterized the conformational dynamics of FL-p53, RNC_1-257_, RNC_1-325_ and RNC_FL_ using HDX-MS. We detected few NC peptides from the proline rich region (PRR) and C-terminal region but obtained good sequence coverage of the TAD (100%) and DBD (65-81%), despite the analytical complexity of the system containing ribosomal proteins (Data S2). The DBD in RNC_1-325_ and RNC_FL_ showed similar levels of deuteration to isolated FL-p53, while the DBD in RNC_1-257_ was deprotected throughout (Fig. 6A-D). The DBD therefore folds into a near-native conformation once fully emerged from the ribosome in RNC_1-325_, consistent with results from limited proteolysis and conformation-specific antibodies (Fig. S3).

**Fig. 6.**
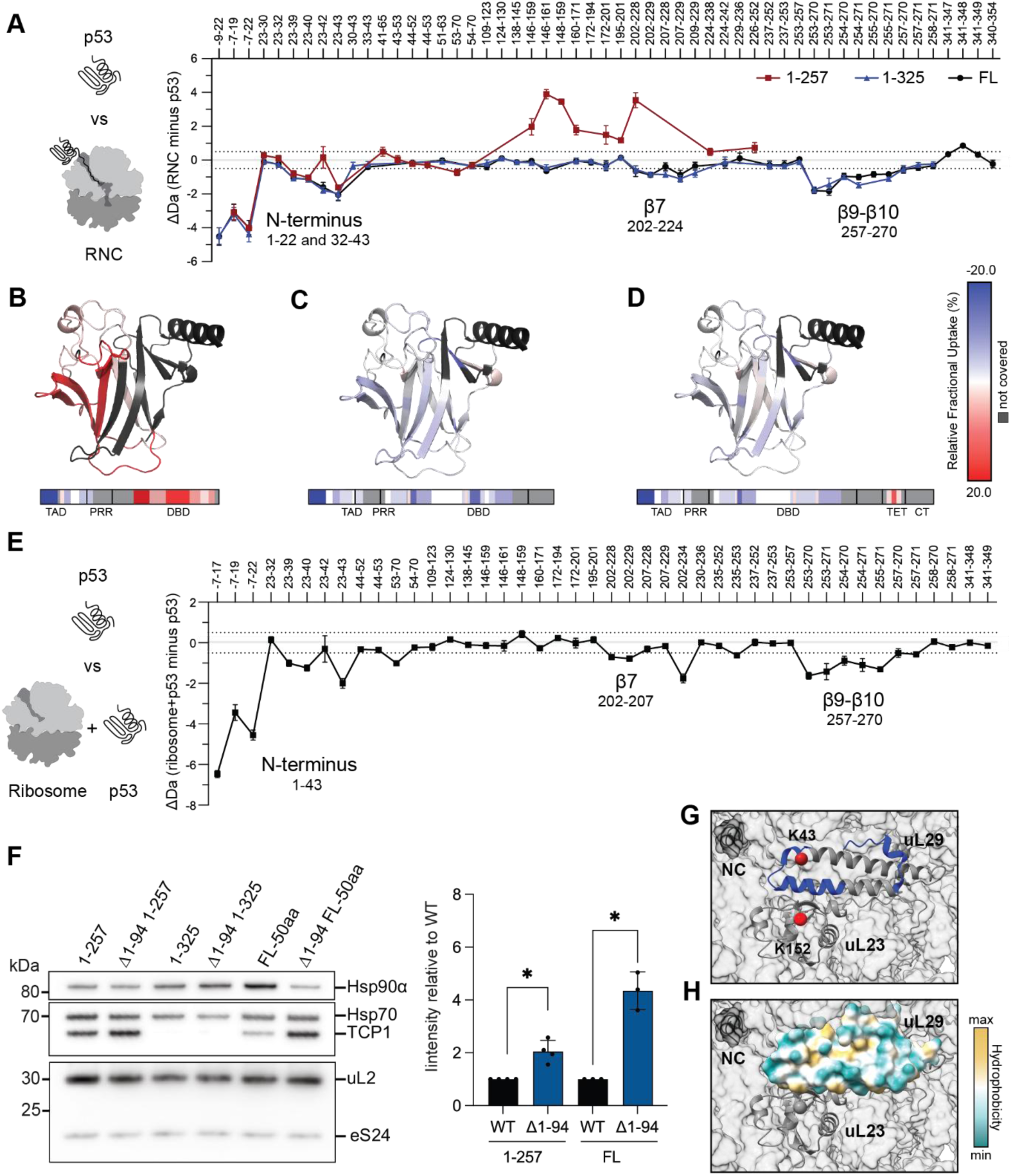
Nascent p53 escapes TRiC by binding the ribosome (E) HDX-MS analysis of p53 NCs. Plot of the mean difference in deuterium uptake (ΔD), after 3 min deuteration, between p53 peptides in RNCs (RNC_1-257_, RNC_1-325_, or RNC_FL_) and isolated full-length p53. Negative values indicate protection of the RNC compared to isolated p53. Dashed lines indicate ±0.5 Da. Data represent mean ± SD, n=3 independent labelling reactions. (**B-D**) Structures of p53 DBD coloured according to the difference in relative fractional uptake, after 3 min deuteration, between isolated full-length p53 and (**B**) RNC_1-257_, (**C**) RNC_1-325_, or (**D**) RNC_FL_. Red indicates increased deuteration of NC peptides relative to isolated p53, and blue indicates decreased deuteration. Regions without peptide coverage are coloured in grey. (E) HDX-MS analysis of isolated p53 mixed with 80S ribosomes. Plot of the mean difference in deuterium uptake (ΔD), after 3 min deuteration, between p53 peptides from a 1:1 mixture of p53 and purified ribosomes, and isolated full-length p53. Negative values indicate protection of p53 upon addition of ribosomes. Dashed lines indicate ±0.5 Da. Data represent mean ± SD, n=3 independent labelling reactions. (F) Removing the p53 N-terminus increases TRiC recruitment to RNCs. Left: WT RNCs and RNCs lacking the N-terminus (Δ1-94) were purified and analyzed by immunoblotting for chaperones. Right: quantification of TCP-1 signal for the indicated RNCs, normalised to the loading control (uL2) and then to WT. Data represent mean ± SD, n=3 biological replicates. Statistical analysis was performed using one-way ANOVA with Dunnett’s post hoc test. Significance relative to control is indicated as p >0.05 (ns), p < 0.05 (*), p < 0.01 (**), p < 0.001 (***), p < 0.0001 (****). (G) Interaction of nascent p53 with the ribosome. Regions of ribosomal protein uL29 that are protected from deuterium exchange in RNC_1-325_ compared to empty ribosomes are coloured blue (PDB: 6OLE). Residues in uL29 and uL23 that crosslink to the NC in RNC_1-325_ are shown as red spheres. (H) As in **(G**), with the surface of uL29 coloured by hydrophobicity. See also Fig S7 and Data S2-S4.

In addition, we noticed that specific regions in nascent p53 were protected relative to isolated FL-p53. Parts of the disordered N-terminal region including the TAD (residues 1-22, 32-43) were protected in all RNCs, and peptides covering β7, β9 and β10 were protected in RNC_1-325_ and RNC_FL_, where the DBD is near natively folded (Fig. 6A-D). To test whether the protection was caused by an interaction with the ribosome, we mixed FL p53 with purified 80S ribosomes and analysed deuterium uptake in p53. This experiment recapitulated the pattern of NC protection we observed for RNCs, indicating that the ribosome binds the folded DBD (Fig. 6E). Importantly, the protected regions included β7, which we showed is important for TRiC recruitment (Fig. 3D). We conclude that ribosome binding buries key elements required for chaperone recognition, explaining why TRiC fails to recognise the ribosome-tethered DBD in RNC_1-325_ and RNC_FL_.

Our data indicated that the p53 N-terminus binds the ribosome in all RNCs, and we wondered whether this also influences chaperone recruitment. Truncating the N-terminus (residues 1-94) did not trigger TRiC binding to RNC_1-325_, indicating that ribosome contacts with β7 suffice to protect the folded DBD from the chaperonin (Fig. 6F). However, the level of TRiC was increased on the Δ1-94 variant of RNC_1-257_, suggesting that the N-terminus interferes with TRiC recruitment during DBD synthesis (Fig. 6F and S7A,B). Mechanistically, the N-terminus may anchor p53 to the ribosome surface (as suggested by HDX-MS) to sterically disfavor TRiC binding. We also considered whether the same phenomenon might regulate cotranslational assembly. Indeed, removing the N-terminus substantially increased the amount of co-assembling p53 on RNC_FL_, with a concomitant increase in TRiC (Fig. 6F and S7C,D). This came at the expense of Hsp90 (Fig. 6F and S7B), consistent with our observation that the chaperones compete for DBD binding (Fig 2D).

To identify the NC interaction sites on the ribosome surface, we analysed deuterium uptake in ribosomal proteins near the exit port. Compared to empty ribosomes, surface-exposed peptides in uL29 were protected in RNC_1-325_ and RNC_FL_, but not RNC_1-257_, consistent with an interaction between the folded DBD and uL29 (Fig. 6G and S7E). As an orthogonal probe of ribosome:NC interactions, we analysed RNC_1-325_ using crosslinking coupled to mass spectrometry (XL-MS). Consistent with HDX-MS, we identified crosslinks between the DBD and both uL29 and the nearby uL23 (Fig. 6G and S7F). The protected surface on uL29 is hydrophilic, consistent with binding to the surface of the folded DBD (Fig. 6H).

In summary, we find that the N-terminus and folded DBD of nascent p53 persistently interact with the ribosome surface near the exit port. Ribosome:NC interactions antagonize both TRiC recruitment and cotranslational assembly until translation termination.

## DISCUSSION

Here, we identify and rationalize the chaperone binding events that direct cotranslational folding of p53, revealing how a model client navigates the cytosolic chaperone network (Fig. 7).

**Fig. 7.**
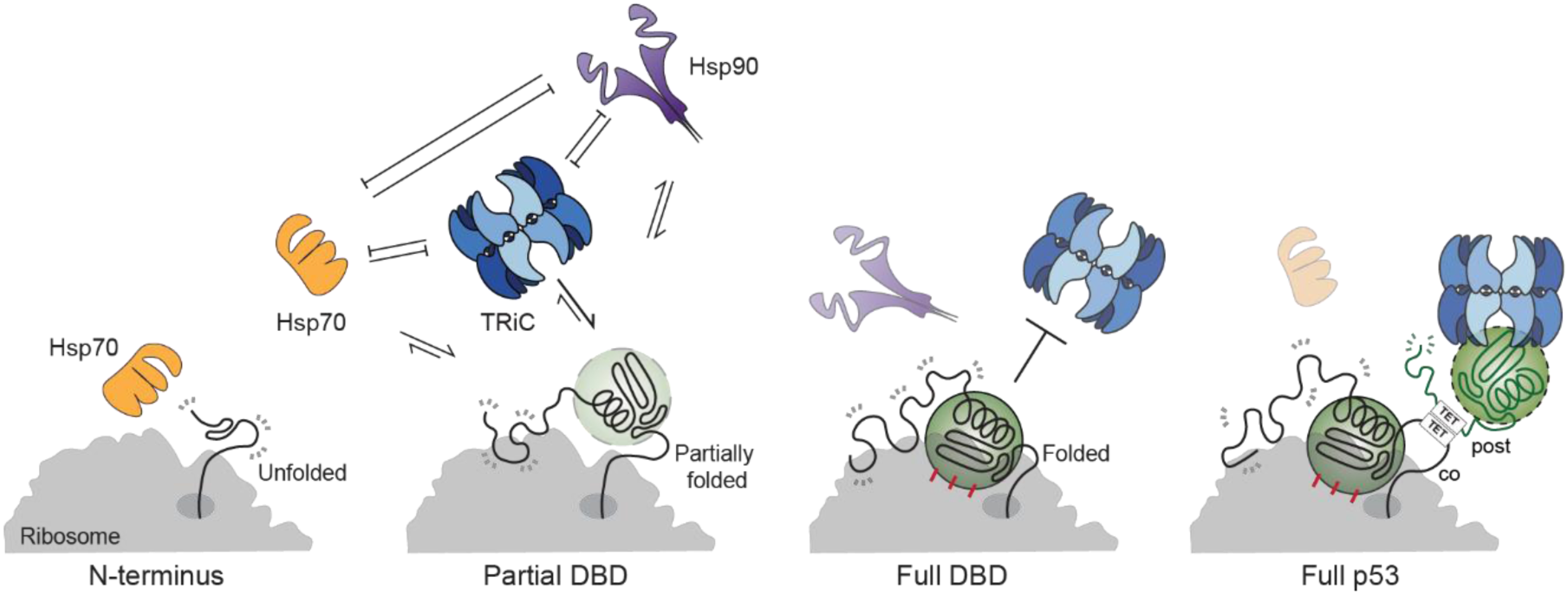
Principles of chaperone prioritization during p53 synthesis Abundant cytosolic chaperones compete to access nascent p53, influenced by local NC folding, NC:ribosome interactions and cotranslational assembly. Hsp70 binding dominates early in synthesis, when only the unstructured N-terminus of p53 is exposed. Subsequent emergence of a folding-incompetent fragment of the DBD recruits TRiC and Hsp90, which compete for binding. Once the DBD is completely synthesised, it folds and excludes TRiC but retains a weak interaction with Hsp90. The exclusion of TRiC is not driven by DBD folding alone, but also by binding of the nascent DBD and p53 N-terminus to the ribosome surface. Towards the end of translation, exposure of the TET domain allows p53 to assemble with an already-synthesized monomer. Since TRiC recognises monomeric p53 post-translation, cotranslational assembly indirectly recruits TRiC to the ribosome.

### Chaperone competition underlies binding hierarchies at the ribosome

Our findings suggest a general framework for understanding cotranslational chaperone action in human cells, wherein constant competition between cytosolic chaperones is tuned by local NC folding and NC:ribosome interactions. Chaperones such as TRiC, Hsp70, Hsp90 and NAC are highly abundant^61^, and we propose that they continuously sample the elongating NC. As translation progresses, subtle changes in NC conformation, exposed sequence or ribosome binding tip the balance between chaperone pathways. In support of this idea, we observe that multiple chaperones bind competitively to nascent p53, and small changes to the NC can alter the chaperone interactome. This behaviour is also consistent with recent measurements of TRiC binding to NCs *in vivo*^18^, which showed that the chaperone frequently and transiently engages translating ribosomes. Considering dynamic NC sampling by chaperones, our data emphasize the importance of quantifying all contemporaneous NC interactors to identify the predominant pathways.

### Structural determinants of chaperone recruitment to NCs

We find that chaperone binding is surprisingly insensitive to perturbation of the global thermodynamic stability of complete ribosome-tethered domains. Conversely, mutations that destabilize the native state can perturb chaperone recruitment during domain synthesis by removing hydrophobic residues that drive chaperone binding. Thus, in contrast to mature proteins, aberrant NCs are distinguished by the exclusion rather than retention of chaperones. As we show here for p53 Y220C, disease-associated mutants can escape chaperone recognition during translation, potentially exacerbating their deleterious effects.

Among the chaperones that recognize nascent p53, TRiC is distinguished by its stable and selective binding. TRiC was previously shown to recognise short sequence-diverse motifs in client proteins^17,62–64^, and our data identify β7/Y220 as one such recognition element in p53. It is likely that several motifs are required for high-affinity binding. Our data further suggest that the conformation of α1/L2 acts as a gatekeeper for TRiC binding motifs. This region is conformationally labile even in the folded DBD^65,66^, potentially explaining why TRiC also binds mature p53 post-translation^30^. Selective binding is consistent with “surgical” chaperone action, with TRiC only recruited by specific client conformations. This contrasts with Hsp70/Hsp90, which extensively unfold then refold mature p53^31,32^.

### Cotranslational assembly

We find that p53 can assemble in a “co-post” fashion on the ribosome, with a mature subunit engaging an NC exposing the TET domain^67^. The alternative “co-co” assembly pathway, involving two NCs, is unlikely. The ribosome exit tunnel occludes ∼30 aa of the NC^68^, and TET is only 37 aa from the C-terminus. TET would therefore be exposed on the leading ribosome at the point of translation termination, but not simultaneously on the trailing ribosome which is at minimum 10 codons behind^69^. Furthermore, disome selective profiling did not detect assembly between p53 NCs in cells^70^. In contrast, a previous study showed that p53 variants did not form mixed dimers when co-expressed in vitro^59^, arguing that p53 NCs assemble on the same polysome in cis. How is this apparent paradox resolved? We propose that co-post assembly of p53 occurs on the same polysome. Upon translation termination, a newly-synthesized p53 monomer is most likely to assemble with a nearby ribosome-tethered NC, thus favouring homomers encoded by the same mRNA. Such a mechanism might be prevalent for homomeric proteins with assembly interfaces towards their C-termini^71^, including *E. coli* βgal as we showed previously^13^. Although we robustly detect cotranslational p53 assembly in cells, we find that assembly is delayed by both TRiC binding and ribosome:NC interactions. Inter-NC interactions may be regulated to avoid misassembly before folding is complete.

### The ribosome controls NC recognition by chaperones

The ribosome is established to act as a binding platform for a subset of chaperones, NC processing enzymes and targeting factors^72–74^. We show that the ribosome plays a more general role in directing chaperone recognition during protein biogenesis. Our finding that TRiC binding is strongly influenced by NC:ribosome interactions may explain a previous observation that TRiC selects distinct clients co-and post-translation^62^. Since not all chaperones are equally affected (i.e. TRiC is excluded but not Hsp70/90), the ribosome acts as an arbiter, dictating the spectrum of chaperones that act during protein synthesis. The ribosome itself may substitute as an NC chaperone. We find that the ribosome buries the highly aggregation-prone β9 in p53^29^, potentially mitigating cotranslational misfolding^75^. Since chaperone binding typically competes with folding^1^, outsourcing chaperone functions to the ribosome may have the advantage of protecting incipient structure in the NC.

### Limitations of this study

Our use of stalled RNCs enables precise control of NC properties, but may bias interactomes compared to ongoing translation. We identify sites on the NC that bind the ribosome, but do not comprehensively identify the parts of the ribosome that bind the NC; XLMS and HDX-MS would not reveal interactions with RNA. Future efforts will need to identify these sites, establish their function, and understand whether the effects we observe for p53 are general.

## RESOURCE AVAILABILITY

### Lead contact

Correspondence and request for materials should be addressed to David Balchin

### Materials availability

Materials are available from David Balchin upon request under a material transfer agreement with The Francis Crick Institute.

### Data and code availability

- All mass spectrometry data have been deposited to the ProteomeXchange Consortium via the PRIDE partner repository with the following dataset identifiers: RNC interactomes: PXD077141

### HDX-MS: PXD077209

- This paper does not report original code
- Any additional information required to reanalyze the data reported in this paper is available from the lead contact upon request.

## ACKNOWLEDGEMENTS

We thank Alžběta Roeselová, David Briggs, Andreas Jörger, Karen Vousden, Maruf Ali and Anne Schreiber for advice, and all members of the Protein Biogenesis laboratory for help and discussion. D.B.’s work is supported by the Francis Crick Institute which receives its core funding from Cancer Research UK (CC2025), the UK Medical Research Council (CC2025), and the Wellcome Trust (CC2025), and by UK Research and Innovation (UKRI) under the UK government’s Horizon Europe funding guarantee [FoldingMap, EP/X020428/1].

## AUTHOR CONTRIBUTIONS

L.K. performed all experiments except MD simulations, and wrote the manuscript together with D.B. T.B.V. performed MD simulations. S.L.M. and J.M.S. analysed crosslinked samples. L.Ke. and J.M.S. collected and partially processed the proteomics data. G.A.P. contributed to optimization of RNC preparation and HDX-MS workflows. D.B. conceived and supervised the project, and wrote the manuscript together with L.K.

## DECLARATION OF INTERESTS

The authors declare no competing interests.

## METHODS

### Experimental Model details

#### Cell lines

Expi293F suspension cells (ThermoScientific) were cultured in FreeStyle 293 Expression Medium (ThermoScientific) at 37°C with 8% CO_2_, under shaking at 125 rpm in vented Erlenmeyer flasks (Corning) and passaged twice at a density of 0.4×10^6^ cells/mL. Cultures were maintained at the following volumes: 30 mL in 125 mL flasks, 50 mL in 250 mL flasks, 100 mL in 500 mL flasks, 200 mL in 1 L flasks, and 600 mL in 2 L flasks. A Vi-CELL XR cell counter (Beckman Coulter) was used to assess cell number and viability.

293T (HPA cultures) cells were maintained in complete DMEM (ThermoScientific) supplemented with 10% FBS (ThermoScientific) and 1% Pen/Strep (ThermoScientific) at 37°C with 5% CO_2_ and passaged at a ratio of 1:10 every 3 days.

## Method details

### DNA vectors and cloning

RNC constructs for human cell expression were synthesised in mammalian expression vector pTwist CMV BG WPRE Neo (Twist Bioscience). The constructs contained an N-terminal 3xFLAG tag, a 3C protease site, NC fragment of interest, and XBP1u+ stalling sequence^5^ followed by a 6xHis tag and stop codon. CMV promoter was used and beta globin (BG) and Woodchuck Hepatitis Virus Post-Transcriptional Regulatory Element (WRPE) were introduced to enhance protein expression. The vectors were amplified using DH5α bacterial strain in LB medium supplemented with 100 μg ml^−1^ ampicillin. p53 was subcloned from WT R72 p53 pCB6+ plasmid (Addgene) using Gibson Assembly Master Mix (NEB), while other full-length proteins were synthesised by Twist Bioscience. RNC truncations and mutations were produced by PCR using Q5 Site-Directed Mutagenesis Kit (NEB) or Phusion High-Fidelity PCR Kit (NEB). Plasmid DNA for transfections was prepared using PureLink HiPure Plasmid Maxiprep Kit (ThermoScientific) from 250 mL cultures.

### Protein purification Human RNC purification

#### RNC purification buffers

RNC lysis buffer contained 50 mM Tris-HCl pH 7.5, 0.5% NP-40, 5 mM MgCl_2_, 25 mM KCl, 10% v/v Glycerol, 200 mM L-arginine, 1 mM Phenylmethanesulfonyl fluoride (PMSF), 1 tablet/50 mL cOmplete, EDTA-free protease inhibitor (Roche), and 2 U/mL RNasin Ribonuclease inhibitor (Promega). RNC sucrose cushion buffer contained 35% w/v RNA/DNAse-free sucrose, 50 mM HEPES-NaOH pH 7.5, 5 mM MgCl_2_, 25 mM (low-salt) or 500 mM (high-salt) KCl, 200 mM L-arginine HCl, and 2 U/mL RNasin Ribonuclease inhibitor (Promega). RNC buffer contained 20 mM HEPES-NaOH pH 7.5, 5 mM MgCl_2_, 25 mM (low-salt) or 500 mM (high-salt) KCl, 200 mM L-arginine HCl, 10 mM NH_4_Cl, 10% v/v glycerol, and 2 U/mL RNasin Ribonuclease inhibitor (Promega). All buffers were made with RNase-free and DNase-free molecular-biology-grade H_2_O.

#### Transient transfection and purification of RNCs

RNCs were transfected and purified as previously described^5^ with modifications to stabilise the interactome. Suspension Expi293F cells were exchanged to fresh FreeStyle 293 Expression Medium (ThermoScientific) and diluted to 3-3.3 x10^6^ cells/mL in 9/10^th^ of the final culture volume. A 3:1 mass ratio of polyethylenimine (PEI):DNA was used for transfections with 1 μg of plasmid DNA per 1 mL of cells to be transfected. PEI (1mg/mL stock, pH 8.0, Polysciences) was diluted in Opti-MEM Reduced Serum Medium with Glutamax (ThermoScientific) to 30 μg/mL in half of the 1/10^th^ final culture volume. Plasmid DNA was resuspended in the same volume of Opti-MEM. The solutions were incubated for 5 min at room temperature and then mixed together, followed by incubation for 20 min at room temperature. The mixture was then added dropwise while swirling the cell culture. The cells were incubated for 16-20h under standard subculture conditions. Cells were harvested by centrifugation at 500 g and washed with ice-cold PBS supplemented with cOmplete EDTA-free protease inhibitor tablet (Roche). For RNC purification, only freshly harvested cells were used. Following wash with PBS, cells were resuspended in RNC lysis buffer at 1/100^th^ cell culture volume and supplemented with RNase-free DNase I (Qiagen) at 10 μL/1 g wet-cell weight. Following 30 min lysis on ice, cell debris was removed by centrifugation at 14,000 g for 15 min at 4°C. The clarified lysate was layered on RNC sucrose cushion buffer and subjected to centrifugation at 70,000 rpm/250,000 g in a Beckmann TLA-110 rotor for 2 h at 4°C to enrich for ribosomes. The ribosome pellets were washed twice and resuspended overnight in RNC buffer at 4 °C with gentle agitation. Resuspended ribosomes were incubated at a ratio of 1 μL of beads per 1 mL of cell culture with anti-Flag M2 Affinity Gel (ThermoScientific), pre-equilibrated with RNC buffer, for 3 h at 4 °C on a rotator. The flow-through was collected by centrifugation (7,000 g) at 4 °C, and the RNC-bound beads were washed five times for 5 min with RNC buffer. The RNCs were eluted with 3xFlag peptide (Peptide Chemistry STP, Francis Crick Institute) diluted to 200 ng/μL in RNC buffer for 1 h at 4 °C on a rotator. Unless indicated otherwise, low-salt sucrose cushion and RNC buffers were used. For HDX-MS and XL-MS, RNCs were purified under a high-salt sucrose cushion and washed with high-salt RNC buffer to remove weakly-bound interactors. The samples were always resuspended and eluted in low-salt RNC buffer.

### Purification of 80S ribosomes

Crude 80S ribosomes were purified from Expi293F cells, following the RNC purification strategy with some differences. During lysis, 2.5 mM puromycin (Santa Cruz Biotechnology) was added to release the NCs from the ribosome. Cell debris was removed as usual, and the ribosomes were isolated from clarified lysate as described above. Ribosome pellets were washed and resuspended overnight. For HDX-MS, high-salt sucrose cushion buffer was used.

### RNC purification with siRNA treatment

293T cells were seeded at 1×10^6^ cells/well in T75 plates (ThermoScientific) and allowed to adhere overnight. The cells were transfected with siRNA (Merck) using Lipofectamine RNAiMax (ThermoScientific) following the manufacturer’s protocol. Briefly, 200 pmol of siRNA and 20 μL of Lipofectamine RNAiMax reagent were separately diluted in Opti-MEM Reduced Serum medium. The Lipofectamine RNAiMax mixture was added to the siRNA and incubated for 10-15 min, and then gently added to the cells to minimise detachment. After 48 h, the cells were transfected with 20 μg DNA using 30 μL Lipofectamine (ThermoScientific) and 40 μL P3000 (ThermoScientific). The DNA/P3000 and lipofectamine solutions were incubated separately at room temperature for 5 min, then combined and incubated for an additional 15 min before gentle addition to the cells. The cells were harvested 72h post siRNA transfection. Harvested cells were lysed, clarified, enriched for ribosomes using a sucrose cushion, and RNCs were immunoprecipitated using anti-Flag M2 Agarose Gel as described above (ThermoScientific). RNCs were eluted with 3xFlag peptide (Peptide Chemistry STP, Francis Crick Institute) diluted to 200 ng/μL in RNC buffer.

The sequences of siRNA duplexes were as follows: TCP1 pool (GGAUGAUAUUGGUGAUGUA, GAAGCAGUGCGUUAUAUCA, GAAGUCAAAUGGAGAGUAU)^30^, CCT5 pool (CUGCUCGUGUUGCUAUUGA, GAAGCAACAGCAUGUCAUA, CCACUUCUGUGAUUAAGUA)^30^, HspA1 (CGACGGAGACAAGCCCAAG)^76^, Hsp90α (GGAAAGAGCUGCAUAUUAA)^77^, control luciferase (CGUACGCGGAAUACUUCGA)^78^. The working stocks (10 μM) were stored at - 20°C for up to 6 months, and the original stocks (100 μM) were maintained at-70°C.

### Immunoprecipitation with conformation-specific antibodies

For conformation-specific antibody experiments, immunoprecipitation with Dynabeads Protein G beads (ThermoScientific) was performed. Briefly, 500 μg of clarified lysate or ribosomal fraction was diluted into lysis/RNC buffer and incubated with the antibody (1:50 dilution) for 1.5 h at 4 °C on a rotator. 30 μL of buffer-equilibrated Dynabeads were added to the sample and incubated for 1.5 h at 4 °C on a rotator. A magnetic rack (Cytiva) was used to isolate and wash the Dynabeads three times. On the last wash, the beads were transferred into a new Eppendorf tube, and the samples were eluted for 10 min at 95 °C in 20 μL Laemmli sample buffer (BioRad).

### Bradford protein assay

Protein concentration was estimated using Protein Assay Dye Reagent Concentrate (BioRad) according to the manufacturer’s instructions. Each sample was mixed with 200 μL Bradford reagent, and the volume was brought up to 1 mL with deionised water. After 5 min incubation, the absorbance at 595 nm was measured. Bovine serum albumin (ThermoScientific) standards were used to generate a standard curve for quantification.

### Immunoblotting

Proteins resolved on SDS-PAGE gels were transferred onto PVDF membranes using the Trans-Blot Turbo System (Bio-Rad). The membranes were blocked in PBS supplemented with 0.1 % w/v Tween 20 (PBS-T) and 5 % w/v non-fat dry milk for 1 h at room temperature. The blots were incubated with relevant primary antibodies at appropriate dilution factors for 1h at room temperature or overnight at 4 °C. After three washes in PBS-T, the membranes were incubated with HRP-conjugated secondary antibodies for 1 h at room temperature or overnight at 4 °C. The membranes were washed with PBS-T three times for 5 min. Membranes were developed using SuperSignal West Pico PLUS Chemiluminescent Substrate (ThermoScientific) and imaged on Amersham Imagequant 800 (Cytiva). When probing for chaperones (TRiC, Hsp70, Hsp90α) and ribosomal proteins (uL2 and eS24), the antibodies were mixed and incubated with the membrane simultaneously, as no cross-reactivity was detected.

### Limited proteolysis

Purified p53 RNCs were diluted to 0.1 μM in RNC buffer, and a reference sample was taken before the treatment with Proteinase K (Millipore) at 2 ng/μL. Samples were incubated at 10°C. Aliquots of reactions were taken at 2.5, 5, 10, and 20 min and quenched with 5 mM pre-chilled PMSF diluted in RNC buffer. The samples were then analysed by SDS-PAGE and immunoblotting.

### Proteomic analysis of RNCs

RNCs and ribosomes were purified under low-salt buffer conditions as described above. 10 μg of purified RNCs or puromycin-treated ribosomes were separated for 8 mm of a 12% 1 mm NuPAGE Bis-Tris SDS-PAGE gel, followed by a Quick Coomassie Stain incubation and destaining with deionised water to remove excess staining. The protein bands were excised and destained in 50% ACN in 50 mM AmBic. After the samples were fully destained, they were reduced for 20 minutes at 56°C with 20mM dithiothreitol (DTT), dehydrated then alkylated for 30 min at room temperature in the dark with 50mM iodoacetamide (IAA). Samples were dehydrated in 100% acetonitrile, digested with 1 μg trypsin overnight at 37°C. Peptides were extracted from the gels with 0.1% TFA followed by 80% ACN in water and eluent spin filtered (Ultrafree-MC centrifugal filters) at 12000 rcf for 4 minutes. Recovered peptides were dried with vacuum centrifugation and then resolubilised using 15 uL of 0.1% TFA and nanodropped for approximate concentration. Digested peptides were loaded onto an UltiMate 3000 HPLC system coupled to a ThermoFisher Scientific Fusion Lumos mass spectrometer using a C18 column (ThermoFisher Scientific PepMap RSLC; 50 cm length, 75 μm inner diameter) operated in a trap and elute configuration. Peptides were eluted over a gradient of 2 % B to 45% B over 45 minutes. (buffer A: 0.1% formic acid, 5% DMSO; B: 80 %(v/v) acetonitrile, 0.1% formic acid, 5% DMSO). Data was acquired in data-dependent acquisition mode. MS1 full scan was performed in the Orbitrap at a resolution of 120,000 from m/z 275-1500. The AGC target was set to standard and the injection time mode was set to auto. The MS2 was performed in the ion trap with the standard AGC target, 20 second dynamic maximum injection time mode, isolation window at 1.2 m/z and 32% normalised HCD collision energy.

Raw data were processed on MaxQuant^79,80^ and Perseus^81^ with UniProt Homo Sapiens reference proteome (UP000005640), and the sequence of FL p53, including the N-terminal FLAG tag and XBP1u+ stalling sequence. A reverse-sequence decoy database was employed to filter false positives, with peptide and protein false discovery rates set at 1%. Proteins were quantified based on iBAQ (intensity-based absolute quantification) values, which were normalised to the mean iBAQ value of 60S ribosomal proteins. The interactome was classified into protein classes based on the Human Proteostasis Network Annotation^82^:

### Equilibrium HDX-MS

RNC labelling, quenching, and offline digestion were performed following the previously published protocol^5,83^. 60S RNCs were purified under high-salt buffer conditions as described above. 60 μL of 0.26-0.5 μM RNCs, 0.5 μM p53 (FL or 1-325 truncation) and 0.5 μM ribosomes were labelled in deuteration buffer (20 mM HEPES pH 7.5, 100 mM KCl, 5 mM magnesium acetate, 1 mM DTT and 0.2 M Arginine–HCl, D_2_O), or H-based buffer which was the same as deuteration buffer but prepared with water instead of D_2_O. To test the effect of ribosomes on p53, ribosomes and p53 were mixed and incubated for 10 minutes at RT before proceeding with the labelling reaction. Given low RNC concentrations, samples were labelled during buffer exchange instead of dilution, using 7K 0.5 mL Zeba spin desalting columns pre-equilibrated with labelling buffers according to the manufacturer’s protocol. Proteins were exchanged and eluted for 2 min at 25 °C, followed by incubation at 25 °C with 350 rpm shaking in an Eppendorf ThermoMixer. The sample was quenched at 3 min on ice by mixing 50 μL of labelled sample with 20 μL quench buffer (175 mM sodium phosphate pH 2.1, 17.5 mM TCEP and 5 M guanidinium chloride, adjusted to pH to 1.2 with orthophosphoric acid) for a final pH^read^ 2.3 after quench. The quenched sample was immediately added to 20 μL of 50% pepsin POROS slurry^84^, which was pre-incubated at 10 °C. The sample was digested offline for 2 min at 10°C with 450 rpm shaking, with quick vortexing every 30 s. After digestion, peptides were eluted through a 0.2 μm PVDF centrifugal filter (Durapore) at 0 °C for 30 s at 16,000 g. The samples were flash frozen and stored at-80 °C prior to analysis.

Samples were thawed and immediately injected into an HPLC coupled to a Synapt G2-Si HDMS^E^ mass spectrometer (Waters) with chromatographic columns kept at 0 ± 0.2 °C during data collection. Digested peptides were trapped and desalted for 4 min at 200 μL/min on a VanGuard pre-column trap (2.1 mm x 5 mm Acquity UPLC BEH C4 1.7 μm (Waters)). Peptides were eluted and separated over a 3-35% acetonitrile in 0.1% formic acid gradient for 25 min at a flow rate of 90 μL/min using a reverse phase Acquity UPLC HSS T3 column (1.8 μm, 1 mm x 50 mm, Waters). Analysis was performed with a Waters Synapt G2-Si in ion-mobility (IMS) mode with a conventional electrospray source and scanning across a 50 to 2000 m/z range. Acquisition parameters were set to a capillary voltage of 3 kV, sampling cone at 40 V, trap collision energy at 4 V, source temperature of 80 °C, and desolvation temperature of 180°C. The instrument was calibrated by direct infusion of [Glu1]-Fibrinopeptide B (50 fmol/μL, Merck) at 5 μL/min. All experiments were carried out under identical conditions without correcting for back-exchange.

Data were analysed as described previously^5,83^. Briefly, peptides were identified from MS^E^ data of undeuterated p53 samples using Protein Lynx Global Server (PLGS 3.0.3, Waters). Peptide masses were identified using information from a nonspecific cleavage database, which contained sequences of p53, all ribosomal proteins, and porcine pepsin. Searches were completed using low energy threshold at 135, elevated energy threshold at 30, and lock mass window at 0.25 kDa. DynamX 3.0 (Waters) was used to process peptides identified by PLGS. The following filters were applied: maximum peptide length of 35 amino acids, minimum product of 0.05 per amino acid, and a minimum of 1 consecutive product identified. Peptides were manually filtered to remove any poor quality or misassigned ribosomal peptides in p53 samples. Additionally, only peptides also identified in p53 off-ribosome control were retained. The relative deuterium uptake (in Da) was determined by subtracting the centroid mass of undeuterated peptides from the matching deuterated peptides. Mean deuterium uptake values are reported as relative measurements, as no correction for back-exchange was performed. Uptake differences over 0.5 Da were considered significant. Data were collected in three technical replicates. All peptide-level data are reported in Data S2 and Data S3.

### Crosslinking MS

RNCs were purified as for HDX-MS, but resuspended and immunoprecipitated in low-salt RNC buffer without arginine and NH_4_Cl after the sucrose cushion. Approximately 400 μL of 0.2 μM RNCs were crosslinked with 1 mM MS-cleavable disuccinimidyl dibutyric urea (DSBU, ThermoScientific) at 25 °C for 1 h with gentle agitation. The crosslinking reaction was quenched with 20 mM Tris-HCl, pH 8.0, for 15 min at 25 °C. Crosslinked samples were processed as described previously^5,84^. Briefly, samples were reduced (10 mM DTT), alkylated (50 mM iodoacetamide), and digested with trypsin. The initial digestion was performed with an enzyme:substrate ratio of 1:100 for 1 h at 25 °C, followed by a second digestion with an enzyme:substrate ratio of 1:20 overnight at 37 °C. Resulting tryptic peptides were separated into five fractions (10%, 20%, 30%, 40%, and 80% acetonitrile in 10 mM ammonium bicarbonate, pH 8.0) using high-pH reverse-phase chromatography on TARGA C18 columns (Nest Group Inc.). Fractions were lyophilised and resuspended in 1% formic acid with 2% acetonitrile. Peptides were analysed by nano-scale capillary LC-MS/MS using Vanquish Neo UPLC (ThermoScientific Dionex) at a flow rate of 250 nL/min, a C18 PepMap Neo nanoViper trapping column (5 μm, 300 μm × 5 mm, ThermoScientific Dionex), an EASY-Spray column (50 cm × 75 μm ID, PepMap C18, 2 μm particles, 100 Å pore size, ThermoScientific), and a quadrupole Orbitrap mass spectrometer (Orbitrap Exploris 480, ThermoScientific). Data acquisition settings were set to data-dependent mode using a top 10 method, beginning with a full high-resolution scan (R=60,000, m/z 380-1,800), followed by a higher-energy collision dissociation of the ten most abundant precursor ions (MS peaks), with ions of 1+ and 2+ charge states excluded.

Data were analysed by first converting Xcalibur raw files to MGF format with Proteowizard^85^. The crosslinked peptide search was performed with MeroX^86^ against a protein database with sequences of 80S ribosome proteins, p53, TRiC, Hsp70s (HSPA1, HSPA8, HSPA6), Hsp90s (Hsp90α, Hsp90β), along with a set of software-generated random decoy sequences. The following parameters were used for the searches: maximum number of missed cleavages of 3, minimum peptide length 5 amino acids, an FDR cut-off of 5%. The variable modifications were set to carbamidomethylation of cysteine (mass shift 57.02146 Da), methionine oxidation (mass shift 15.99491 Da), DSBU modified fragments: 85.05276 Da and 111.03203 Da (precision: 5 ppm MS and 10 ppm MS/MS). Crosslink analysis was performed with XiView^87^. Crosslinks with a score below 50 were excluded from the analysis.

### Molecular dynamics

The DNA-binding domain (DBD, residues 94 to 292) of wild-type p53, p53^R175H^, p53^Y220C^, p53^R248Q^, p53^Δβ3^ and p53^Δβ5^ were modelled by AlphaFold3^88^ before being placed in a 10 × 10 × 10 nm box and solvated in TIP3P water^89^ and 150 mM NaCl (with extra neutralising ions where appropriate). For each p53 variant, three independent simulations were performed in Gromacs 2023.1^90,91^ with the CHARMM36m forcefield^92^. The system was minimised with the steepest-descent algorithm to convergence or a maximum force of 1000 kJ/mol before equilibration in the NVT ensemble at 310K maintained by a v-rescale thermostat (τ_T_ = 0.1 ps) for 100 ps (2 fs time step), followed by further equilibration in the NPT ensemble at 310K (v-rescale thermostat, τ_T_ = 0.1 ps) and 1 bar (isotropic Parrinello-Rahman barostat, τ_P_ = 2.0 ps) for 1 ns (2 fs time step). Production runs were performed for 1 µs (2 fs time step) at 310K (v-rescale thermostat, τ_T_ = 1.0 ps) and 1 bar (isotropic c-rescale barostat, τ_P_ = 5.0 ps). The Verlet scheme was used to impose a cut-off distance of 1.2 nm for both van der Waals and coulombic interactions and the LINCS algorithm^93^ was used to restrain hydrogen bonds. The first 200 ns of each production run were excluded from the analysis to ensure the system had reached equilibrium. The radius of gyration, surface-accessible solvent area and per-residue root-mean squared fluctuation were calculated over the course of the simulations with the built-in Gromacs tools *gmx gyrate*, *gmx sasa*, and *gmx rms* respectively. Plots were generated with seaborn^94^ and Matplotlib^95^ in Python 3.13.2.

## SUPPLEMENTAL INFORMATION

**Data S1.** RNC interactomes

**Data S2.** HDX-MS analysis of p53 RNCs

**Data S3.** HDX-MS analysis of uL29

**Data S4.** XLMS analysis RNC_1-325_

## SUPPLEMENTAL FIGURES AND LEGENDS

**Figure S1.**
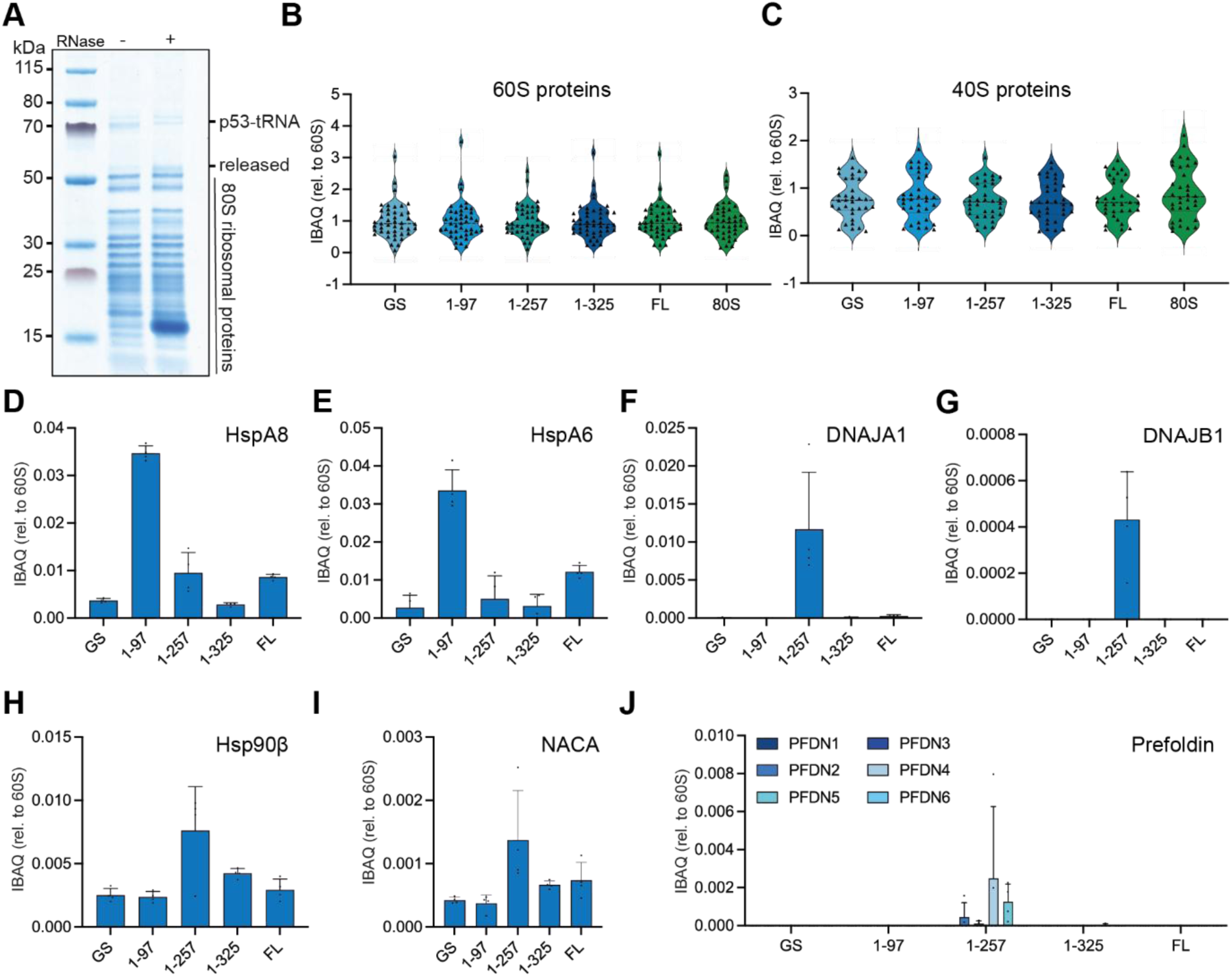
RNC composition and interactors, related to Figure 1 (**A**) RNC integrity. Coomassie-stained SDS-PAGE analysis of p53 RNC_1-325_. The band corresponding to the nascent chain (p53-tRNA) migrates higher due to a covalently linked peptidyl-tRNA, which is degraded by addition of RNase, causing a ∼20kDa shift (released). (**B-C**) RNCs contain intact 80S ribosomes. Mean intensity-based absolute quantification (iBAQ) values for (**B**) 60S ribosomal proteins and (**C**) 40S ribosomal proteins in each RNC. Values are normalised to the average iBAQ of 60S ribosomal proteins in each sample. n=4 biological replicates. (**D-J**) Quantification of chaperone occupancy on RNCs. Mean iBAQ values for (**D**) HspA8, (**E**) HspA6, (**F**) DNAJA1, (**G**) DNAJB1 (**H**) Hsp90β, (**I**) NACA, and (**J**) prefoldin subunits (PFDN1-6). Values were normalised to the average iBAQ of 60S ribosomal proteins in each sample. Data represent mean ± SD, n=4 biological replicates.

**Figure S2.**
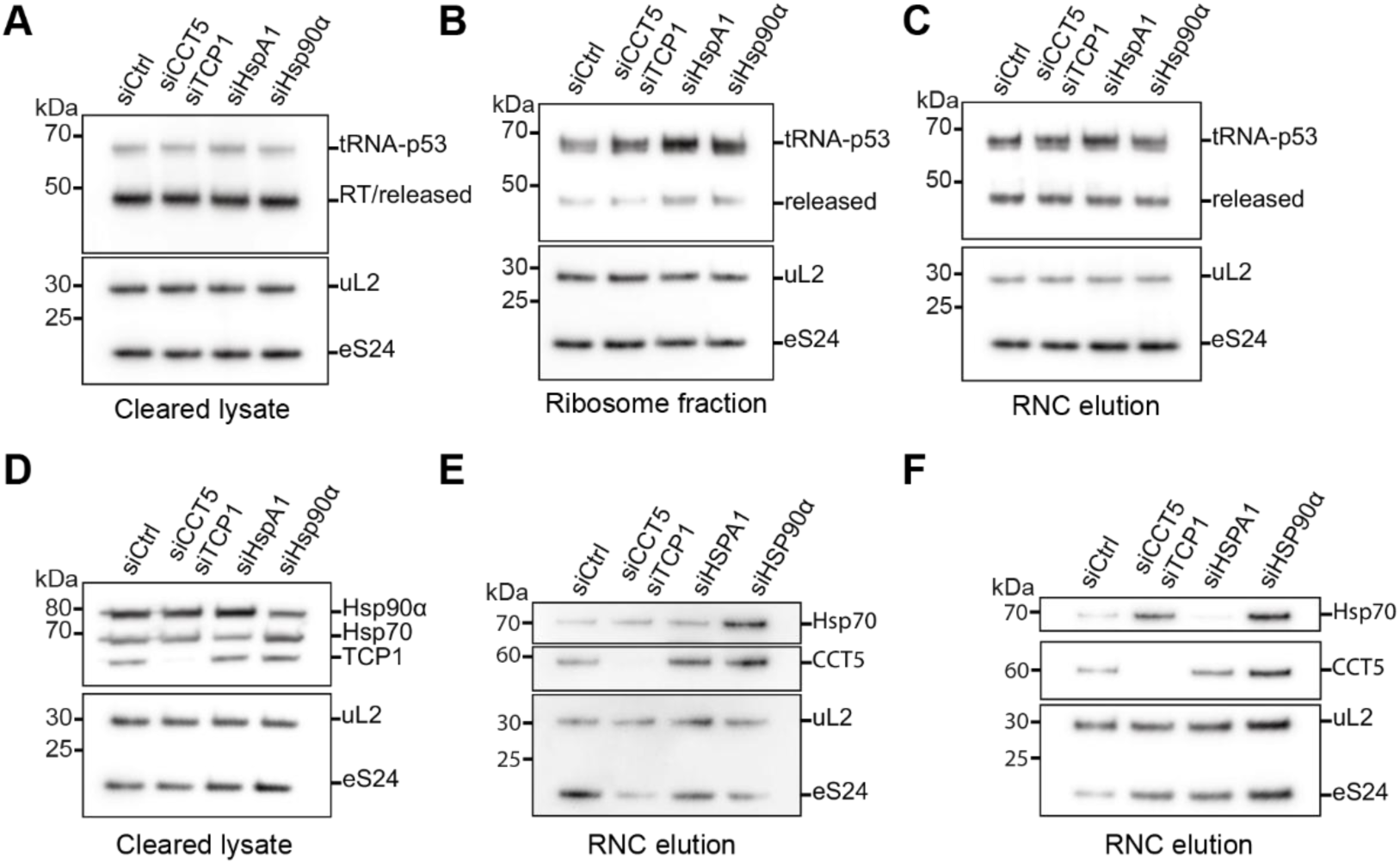
Effect of chaperone knockdown on p53 RNC interactome, related to Figure 2 (**A-C**) Chaperone knockdown does not affect RNC integrity. Cells were treated with control siRNA (siCtrl), or siRNA targeting TRiC subunits (TCP1/CCT5), Hsp70 (HspA1), or Hsp90α, before expression of RNC_1-257_. Anti-FLAG immunoblots of the (**A**) cleared lysate, (**B**) ribosome fraction, and (**C**) purified RNC_1-257_ are shown. Immunoblots were performed for large (uL2) and small (eS24) subunit ribosomal proteins, as controls. (**D**) Chaperone knockdown efficiency. As in (**A**), except that samples were probed for chaperones as indicated. (**E-F**) Effect of chaperone knockdown on RNC interactome, replicates of Fig. 2D. Cells were treated with control siRNA (siCtrl), or siRNA targeting TRiC subunits (TCP1/CCT5), Hsp70 (HspA1), or Hsp90α, before expression and purification of RNC_1-257_. Samples were analyzed by immunoblotting for chaperones as indicated.

**Figure S3.**
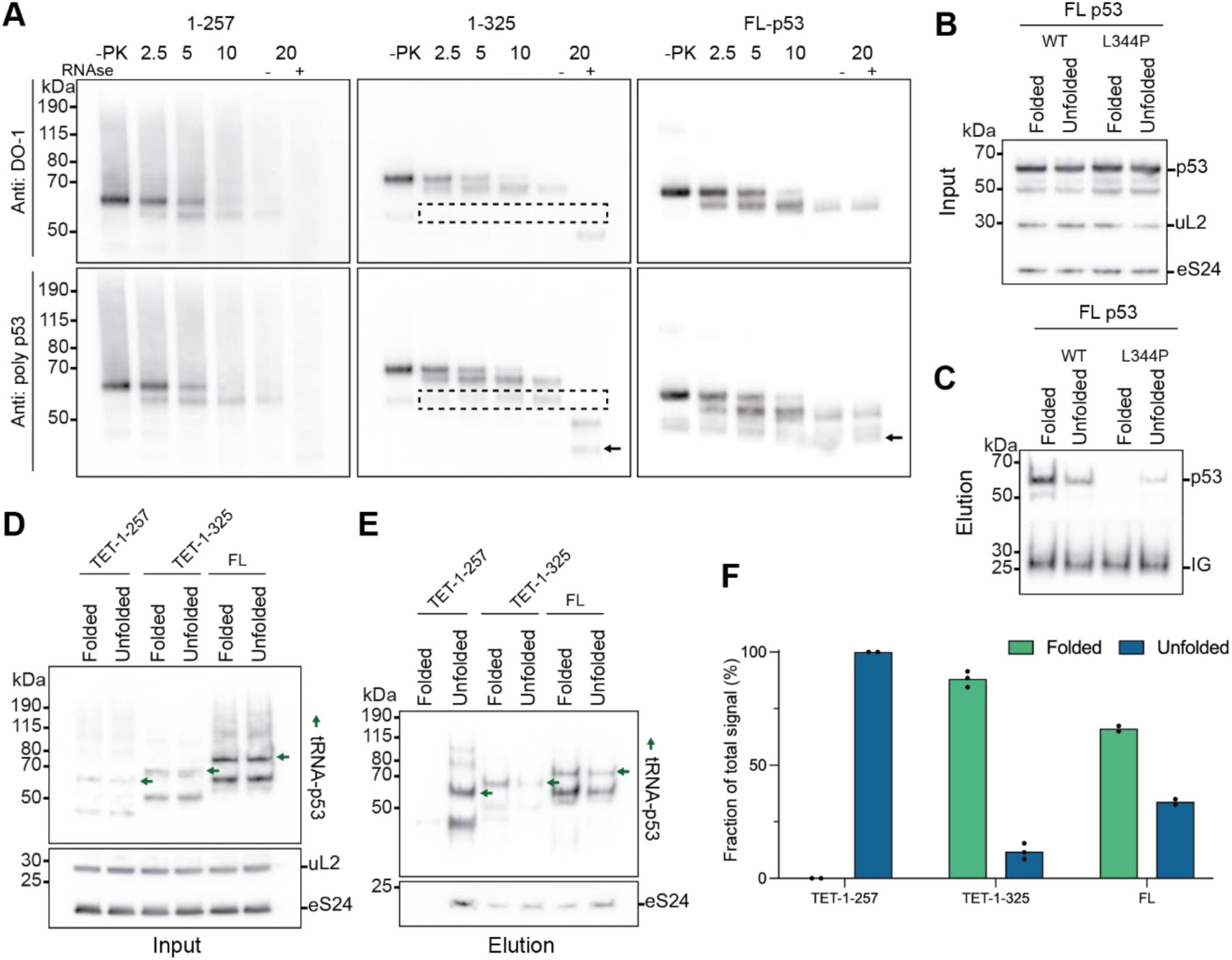
Cotranslational folding of p53 DBD, related to Figure 3 (**A**) Limited proteolysis of RNCs and isolated full-length p53 (FL p53). RNC_1-257_, RNC_1-325_, and FL p53 were treated with 2 ng/μL Proteinase K for 2.5, 5, 10, or 20 minutes. Immunoblots were probed using a monoclonal antibody against an epitope near the N-terminus of p53 (DO-1, epitope aa20-26, top) or a polyclonal p53 antibody (bottom). The box indicates the PK-resistant intermediate in RNC_1-325_ and the arrow indicates the ∼35kDa intermediate observed in RNC_1-325_ (after RNase treatment) and FL p53. (**B-C**) Pab1620 does not immunoprecipitate monomeric p53. FLAG-tagged WT or L344P monomer mutant FL p53 were expressed in Expi293F cells and immunoprecipitated using conformation-specific antibodies: “folded” (Pab1620) or “Unfolded” (Pab240). (**B**) Input and (**C**) elution were analyzed by immunoblotting for FLAG. (**D-E**) Analysis of RNCs using conformation-specific antibodies. FLAG-tagged RNC_TET-1-257_, RNC_TET-1-325_ and RNCwere expressed in Expi293F cells and the ribosomal fraction was isolated before immunoprecipitation using p53 antibodies specific to the “folded” (Pab1620) or “unfolded” (Pab240) conformation. (**D**) Input and (**E**) elution were analyzed by immunoblotting for FLAG. (**F**) Quantification of tRNA-p53 signal from (**E**), expressed as a percentage of total Folded plus Unfolded signal (n=3 for RNC, n=2 for RNCand RNC).

**Figure S4.**
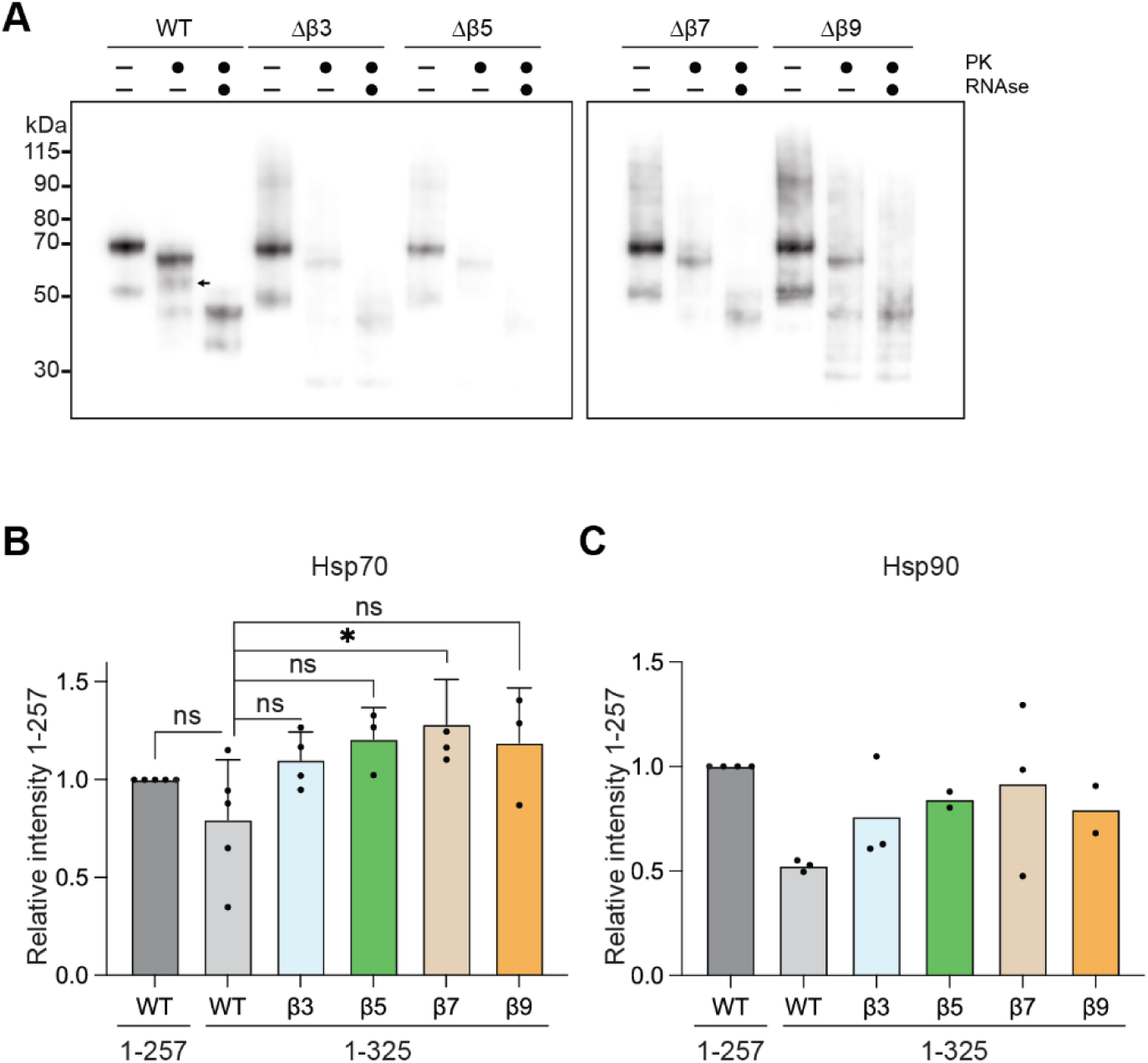
Effects of β-strand deletions on RNC folding and chaperone interactome, related to. **Figure 3** (A) Limited proteolysis of RNCs shows that β-strand deletions are destabilizing. Variants of RNCwere treated with 2 ng/μL Proteinase K for 10 minutes and analyzed by immunoblotting using a polyclonal p53 antibody. Samples were also treated with RNase to identify the PK-resistant species, indicated by the arrow. (B) Hsp70 recruitment to RNCs with β-strand deletions. Quantification of Hsp70 signal from Fig. 3C. Data represent mean ± SD, n=3-5 biological replicates. Statistical analysis was performed using one-way ANOVA with Dunnett’s post hoc test. Significance relative to control is indicated as p >0.05 (ns), p < 0.05 (*). (C) As in (**B**), for Hsp90α. n=2-4 biological replicates.

**Figure S5.**
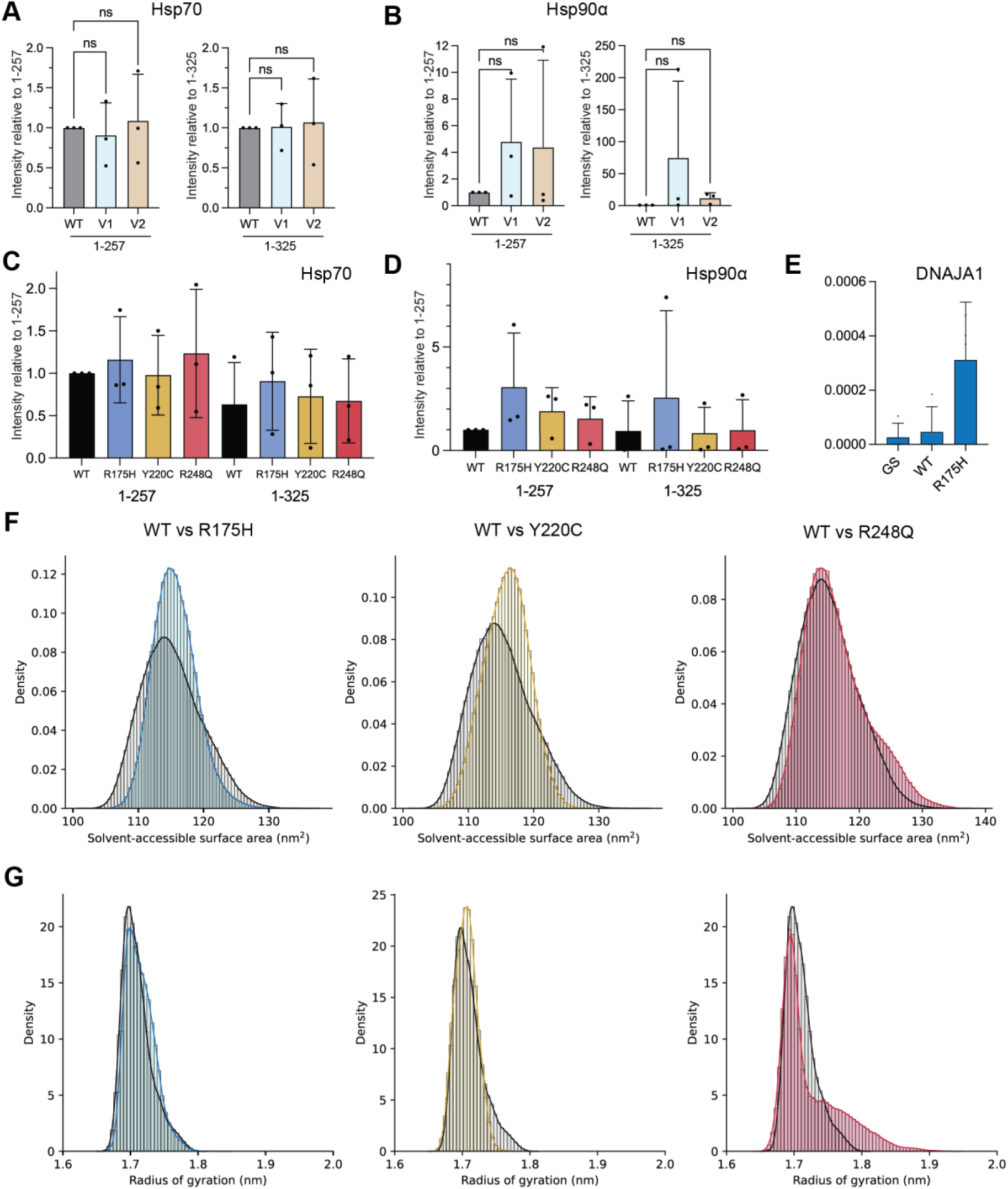
Effects of RNC mutation on NC interactors and conformation, related to Figure 4 (A) Hsp70 recruitment to RNCs with combinations of mutations. Quantification of Hsp70 signal from Fig. 4B, normalised to the loading control (uL2) and then to WT. Data represent mean ± SD, n=3 biological replicates. Statistical analysis was performed using one-way ANOVA with Dunnett’s post hoc test. Significance relative to control is indicated as p >0.05 (ns). (B) As in (**A**), for Hsp90α. n=3 biological replicates (C) Hsp70 recruitment to RNCs with point mutations. Quantification of Hsp70 signal from Fig. 4C, normalised to the loading control (uL2) and then to WT. Data represent mean ± SD, n=3 biological replicates. Statistical analysis was performed using one-way ANOVA with Dunnett’s post hoc test. Significance relative to control is indicated as p >0.05 (ns). (D) As in (**C**), for Hsp90α. n=3 biological replicates. (E) R175H mutation recruits DNAJA1 to RNC. Proteomic analysis of RNCs yielded mean iBAQ values for DNAJA1, which were normalised to the average iBAQ of 60S ribosomal proteins in each sample. n=4 biological replicates. (F) Density plot of the solvent-accessible surface area (SASA) of each p53 DBD single-point mutant (coloured lines and bars) compared to wild-type (black line, grey bars). (G) Density plot of the radius of gyration of each p53 DBD single-point mutant (coloured lines and bars) compared to wild-type (black line, grey bars).

**Figure S6.**
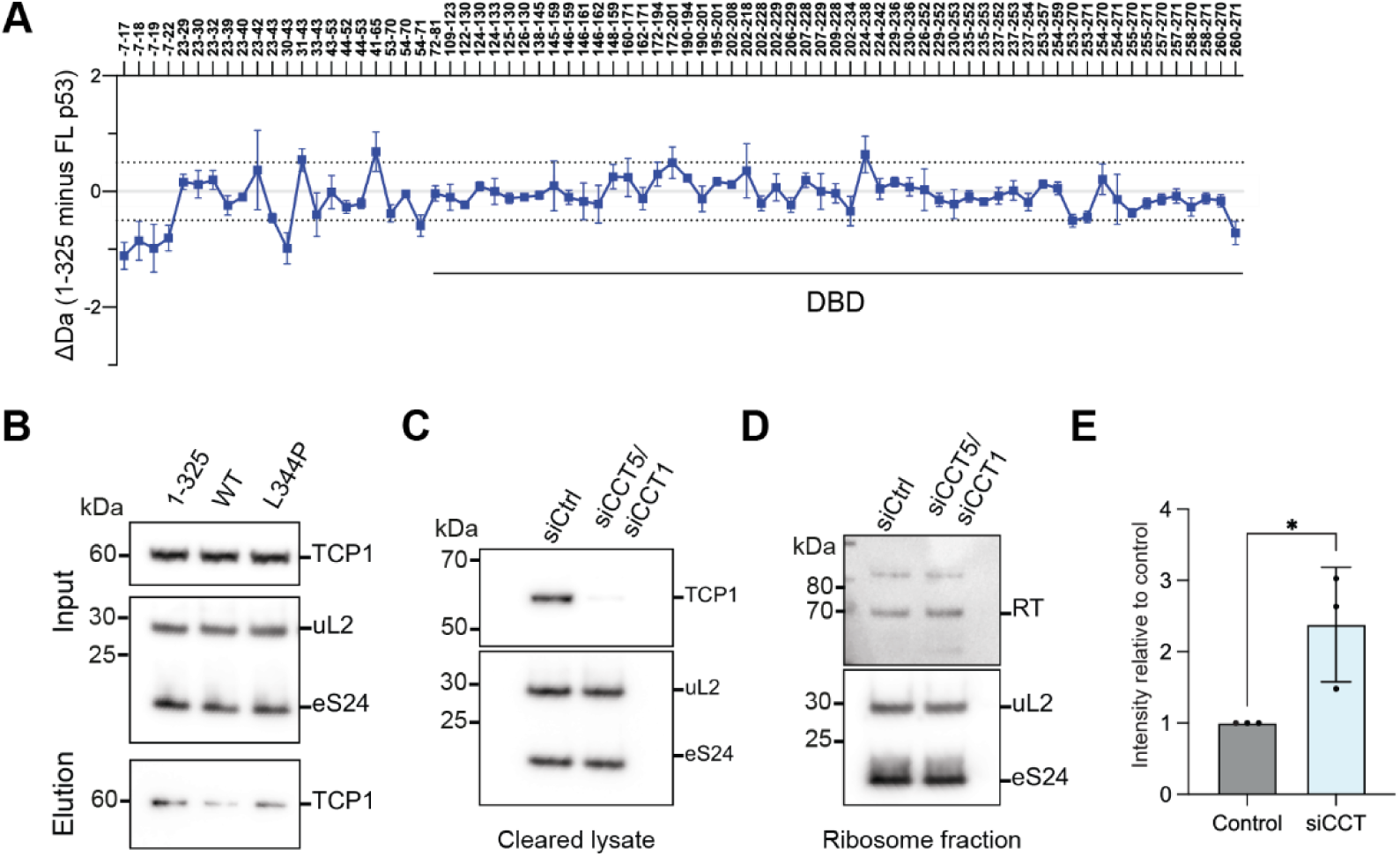
Cotranslational assembly of p53, related to Figure 5 (A) HDX-MS shows that C-terminal regions do not allosterically destabilise the p53 DBD. Plot of the mean difference in deuterium uptake (ΔD), after 3 min deuteration, between p53 1-325 and FL p53, both off the ribosome. Dashed lines indicate ±0.5 Da. Data represent mean ± SD, n=3 independent labelling reactions. (B) TRiC binds monomeric p53 off the ribosome. FLAG-tagged p53 variants were expressed in Expi293F cells, isolated via FLAG pulldown and analyzed by immunoblotting for TRiC (TCP-1). (C) TRiC knockdown efficiency. Cells expressing RNCwere treated with control siRNA (siCtrl), or siRNA targeting TRiC subunits (TCP1/CCT5), and the cleared lysate was analyzed by immunoblotting for TCP1. (D) Cotranslational assembly is enhanced upon TRiC knockdown. Following RNCexpression and siRNA treatment as in (**C**), the ribosomal fraction was isolated and analyzed by immunoblotting for poly-His, indicative of recruitment of mature (RT) p53. (E) Quantification of RT (His) signal from (**D**), normalised to the loading control (uL2) and then to siCtrl. Data represent mean ± SD, n=3 biological replicates. Statistical analysis was performed using unpaired t-test. Significance relative to control is indicated as p >0.05 (ns), p < 0.05 (*).

**Figure S7.**
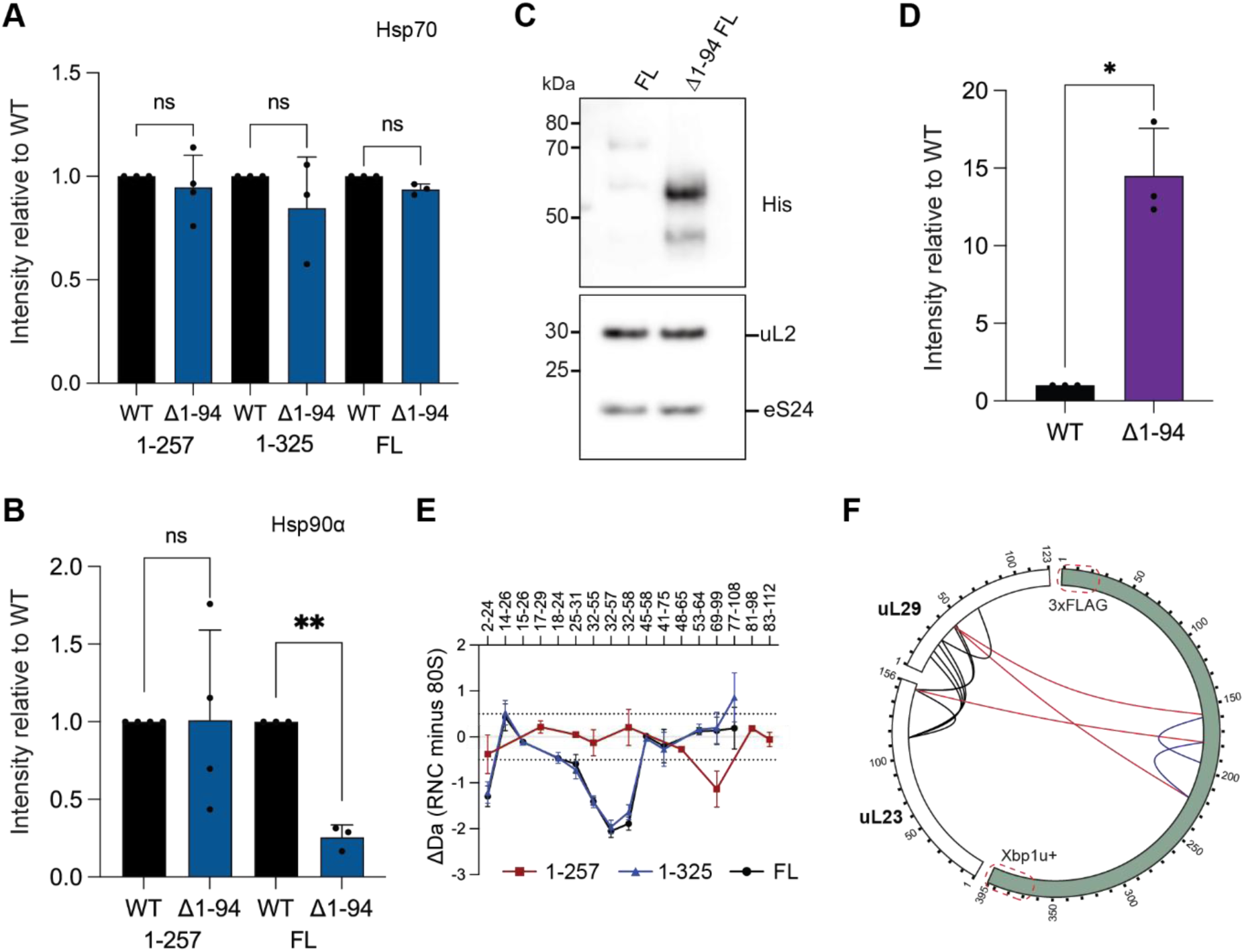
Ribosome interaction with nascent p53, related to Figure 6 (A) Hsp70 recruitment upon deletion of the p53 N-terminal region. Quantification of Hsp70 signal from Fig. 6F, normalised to the loading control (uL2) and then to WT. Data represent mean ± SD, n=3 biological replicates. Statistical analysis was performed using one-way ANOVA with Dunnett’s post hoc test. Significance relative to control is indicated as p >0.05 (ns). (B) As in (**A**), for Hsp90α. Data represent mean ± SD, n=3-4 biological replicates. Significance relative to control is indicated as p >0.05 (ns), p < 0.05 (*), p < 0.01 (**). (C) Deletion of the N-terminus increases cotranslational assembly. WT and Δ1-94 RNCwere purified and analyzed by immunoblotting for poly-His, indicative of recruitment of mature p53. (D) Quantification of His signal from (C), normalised to the loading control (uL2) and then to WT. Data represent mean ± SD, n=3 biological replicates. Statistical analysis was performed using unpaired Welch’s t-test. Significance relative to control is indicated as p >0.05 (ns), p < 0.05 (*). (E) Protection of uL29 by nascent p53. Plot of the mean difference in deuterium uptake (ΔD), after 3 min deuteration, between ribosomal protein uL29 in the indicated p53 RNCs and empty 80S ribosomes. Dashed lines indicate ±0.5 Da. Data represent mean ± SD, n=3 independent labelling reactions. (F) NC crosslinks to uL29 and uL23. Map of DSBU crosslinks in RNCidentified by mass spectrometry. Only proteins that crosslink to the NC are shown.

## Notes

### Competing Interest Statement

The authors have declared no competing interest.

